# Computational design of conformation-biasing mutations to alter protein functions

**DOI:** 10.1101/2025.05.03.652001

**Authors:** Peter E. Cavanagh, Andrew G. Xue, Shizhong Dai, Albert Qiang, Tsutomu Matsui, Alice Y. Ting

## Abstract

Most natural proteins alternate between distinct conformational states, each associated with specific functions. Intentional manipulation of conformational equilibria could lead to improved or altered protein properties. Here we develop Conformational Biasing (CB), a rapid and streamlined computational method that utilizes contrastive scoring by inverse folding models to predict variants biased towards desired conformational states. We validated CB across seven diverse deep mutational scanning datasets, successfully predicting variants of K-Ras, SARS-CoV-2 spike, β2 adrenergic receptor, and Src kinase with improved conformation-specific functions including enhanced effector binding or enzymatic activity. Furthermore, applying CB to lipoic acid ligase, a conformation-switching bacterial enzyme that has been used for the development of protein labeling technologies, revealed a previously unknown mechanism for conformational gating of sequence-specificity. Variants biased toward the “open” conformation were highly promiscuous, while “closed” conformation-biased variants were even more specific than wild-type, enhancing the utility of LplA for site-specific protein labeling with fluorophores in living cells. The speed, simplicity, and versatility of CB (available at: https://github.com/alicetinglab/ConformationalBiasing/) suggest that it may be broadly applicable for understanding and engineering protein conformational dynamics, with implications for basic research, biotechnology, and medicine.

## Introduction

Computational protein design methods have excelled at the design of static protein structures, but the vast majority of natural proteins switch between two or more conformational states in order to carry out their functions. For example, G proteins signal to effectors from their GTP-bound conformation but not their GDP-bound conformation, membrane receptors change conformation upon ligand binding, and many enzymes alternate between distinct inactive and active conformations. As more alternative conformations for proteins of interest are elucidated by structural methods as well as computational tools, the ability to deliberately bias proteins towards one conformational state over another could enable targeted improvements in protein functions. For example, point mutations that shift G proteins towards their GTP-bound conformations could produce stronger or more sustained downstream signaling; a GPCR variant biased towards its active conformation could produce more potent responses to agonists; and variants that stabilize the pre-fusion conformation of viral membrane proteins could produce better vaccines for eliciting neutralizing antibodies.

In a simple example in which a protein of interest (POI) alternates between two conformational states (**Figure 1A**), our goal is to predict point mutations that shift the relative occupancy of those states (**Figure 1B**). Ideally, our method should be extremely fast and scalable (able to characterize thousands of variants in under a minute), generalizable to a wide variety of proteins and protein complexes without extensive tailoring, and accessible to non-specialist labs with limited computational resources. Among existing methods, molecular dynamics simulations(1-5) and physics-based deep-learning models(6) can address this challenge to some extent, but as they are difficult to implement and computationally expensive, they cannot currently be scaled for the routine analysis of thousands of protein variants. AlphaFold is also computationally costly to perform at scale, and has been shown to have limited sensitivity to mutations (7). Recently, AFCluster, which refines AlphaFold predictions through subsampling of protein MSAs (mutiple sequence alignments), was used to predict natural protein variants that exist in alternative conformations. However, AFCluster cannot address non-natural variants or proteins with a shallow MSA, and so far has only been applied to fold-switching proteins (8).

**Figure 1.**
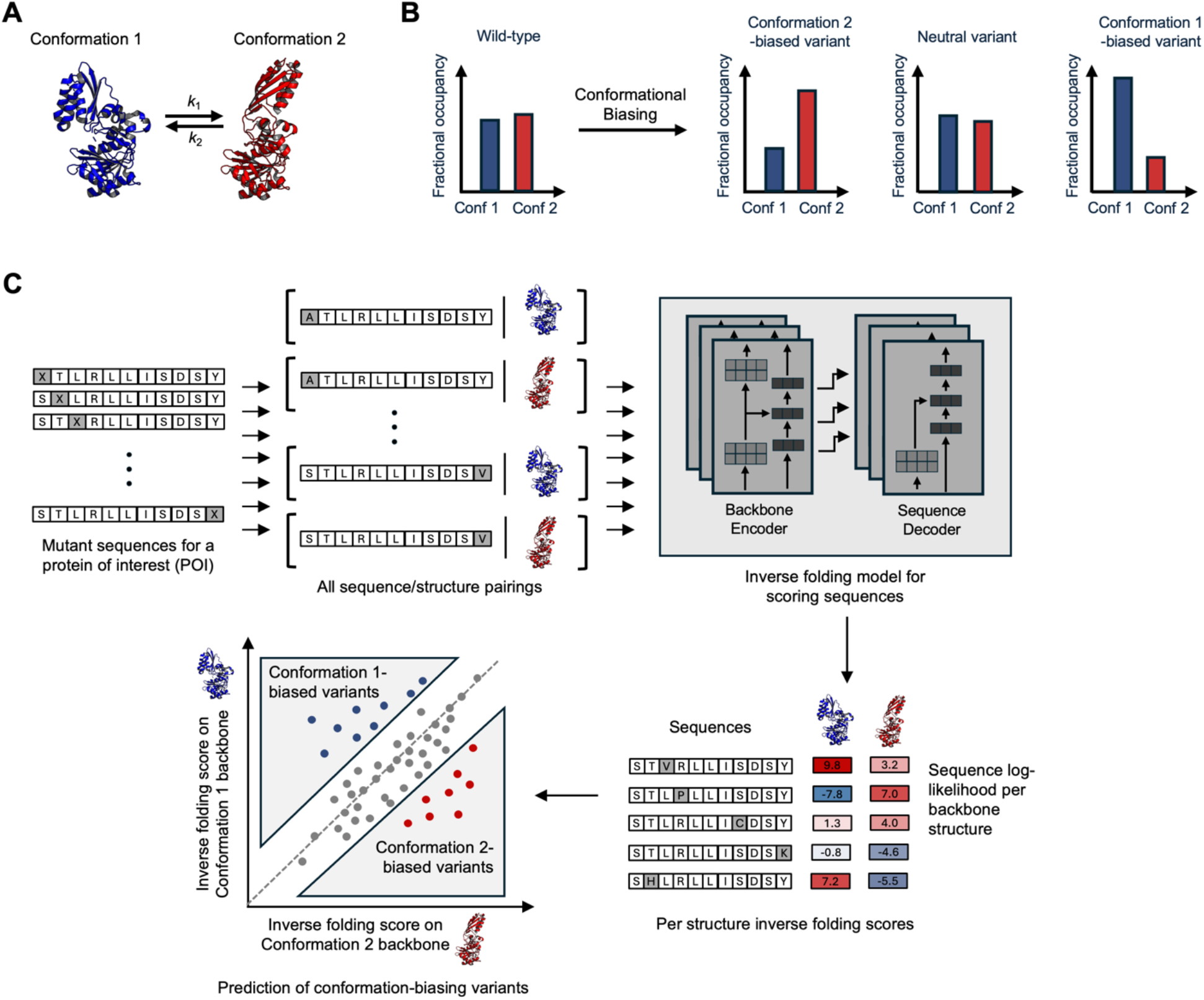
Conformational biasing (CB) approach for predicting mutations that shift conformational equilibria. (**A**) Many proteins switch between alternative conformations. Specific conformations may exhibit unique binding, catalytic, and functional properties. (**B**) CB predicts mutations that shift conformational equilibria. (**C**) Computational workflow for CB. Mutant sequences for a protein of interest are paired with the protein’s structure in two or more conformational states. The structures can be experimentally-solved or computationally-predicted. The sequence-structure pairings are scored using an inverse folding model such as ProteinMPNN and plotted as shown. Conformation-biasing mutations maximize the difference in conformation-specific inverse folding scores.

De novo proteins that exist in multiple conformational states have been generated by sampling for sequences that can exist in all required conformations(9, 10). In some of these studies, inverse folding models (IFMs) and/or AlphaFold have been used to compare sequences designed for different conformations, and in the process uncover positions that may be important for conformational regulation (11-14). IFMs like ProteinMPNN(15) and ESM-IF1(16) are trained to discriminate between structurally similar amino acids at a given position in order to recover wild-type sequence based solely on a protein’s backbone fold. They are therefore especially sensitive to point mutations, a feature that has been leveraged for the design of structure- and complex-stabilizing mutations (17-19). We hypothesized that, despite the subtle conformational changes present in many natural proteins, IFMs could potentially be applied for the *in silico* screening of variant libraries, and help to identify mutations that preferentially stabilize one conformational state over another.

Here we develop a computational workflow called “Conformational Biasing” (CB) for the fast and scalable prediction of mutations that bias proteins toward desired conformational states. CB uses an inverse folding model to generate sequence- and structure-conditioned scores for a library of variants (typically point mutations) of a protein of interest, against two or more alternative backbone structures (**Figure 1C**). Highly divergent scores predict conformational biasing. Surprisingly, we find that no additional fine-tuning, added prediction heads, or custom-training objectives are required to achieve strong performance across multiple datasets for the identification of conformationally-biasing mutations.

We first validated CB using three deep mutational scanning (DMS) datasets of proteins that engage with effectors in a highly conformation-specific manner: the G protein K-Ras; the SARS-CoV2 spike protein which binds to the ACE2 receptor in the “up” conformation but not “down” conformation; and the β2 adrenergic receptor that signals to G proteins and arrestin in its active conformation. We then analyzed four enzyme DMS datasets (Src, B-Raf, FabZ, MurA), to see if CB could predict point mutations that enhance enzyme activity through biasing towards active vs. inactive conformations. Finally, we applied CB to precisely engineer lipoic acid ligase (LplA), a bacterial enzyme(20, 21) that has been harnessed for the ligation of chemical probes to specific proteins in living cells(22). We discovered that biasing LplA towards its “closed” adenylate-ester bound structure improved its sequence-specificity, while variants biased towards the “open” conformation were vastly more promiscuous, capable of tagging endogenous proteomes and laying the groundwork for improved “proximity labeling” technology for unbiased mapping of spatial proteomes. Overall, we find that engineering towards conformation-specific protein behavior can result in larger and more specific improvements in protein function than previous methods.

## Results

We developed the CB workflow shown in **Figure 1C**, which pairs every possible point mutant of a protein of interest (POI) with two (or more) alternative backbone structures. An IFM such as ProteinMPNN(15) is used to score every sequence:structure pairing in a sequence- and structure-conditioned manner, such that the fitness of a mutation as well as its impact on the fitness of residues at other positions is considered. Generally, IFM-predicted beneficial/deleterious mutations are highly correlated across alternative backbone structures. Rather than focusing on overall highest-scored variants as in previous studies using IFM-based scoring(18), however, we specifically turned our attention to variants with the most *contrasting* scores between structures. For example, in the scatter plot in **Figure 1C**, the diagonal represents mutations with high agreement in predicted fitness between structures. Variants below the diagonal, with significantly higher Conformation 2 score than Conformation 1 score are predicted to be “Conformation 2-biased” whereas variants above the diagonal are predicted to be biased toward Conformation 1.

To evaluate CB’s predictions against experimental data, we searched for deep mutational scanning (DMS) studies of proteins known to undergo conformational change. Though we could not find large datasets with direct measurements of protein conformation by NMR, SAXS, or Cryo-EM, we did find several studies using indirect readouts of protein conformation, such as state-specific binding to protein effectors. We reasoned that such datasets might present a challenging evaluation task for CB given the noisiness of biological assays.

We first examined K-Ras, an important G protein that signals to downstream effectors through a GTP-bound State 2 conformation but not GDP-bound State 1 conformation (**Figure 2A**). Oncogenic mutations in K-Ras can alter the State 1: State 2 equilibrium, leading to uncontrolled cell proliferation(23). Weng et al.(24) experimentally evaluated >30,000 single and double K-Ras mutants for binding to a panel of State 1 and State 2-specific proteins. We used CB to predict State 1 or State 2-biased K-Ras mutations. In **Figure 2B**, we show variants plotted by their State 1 and State 2 scores, and colored by experimentally-determined binding to RALGDS, a State 2-specific effector, or DARPin K27, a State 1-specific synthetic binder(33, 38). To exclude variants that act through direct alteration of the effector binding interface, we omitted interface residues from our analysis based on prior structures of K-Ras:effector complexes (**Figure S1A**). **Figures 2B-C** show that State 2-biased mutants predicted by CB exhibit significantly higher RALGDS binding. We performed a similar analysis of binding data for five other State 2-specific effectors reported in Weng et al.(24) and observed, in each case, a consistent and highly significant (p < 0.0001) increase in binding scores for CB-predicted State 2-biased versus State 1-biased K-Ras variants (**Figure 2C**). In contrast, CB-predicted State-2 biased variants exhibit the opposite effect on DARPin K27 binding (which is State 1-specific), with significantly reduced binding compared to State 1-biased variants.

**Figure 2.**
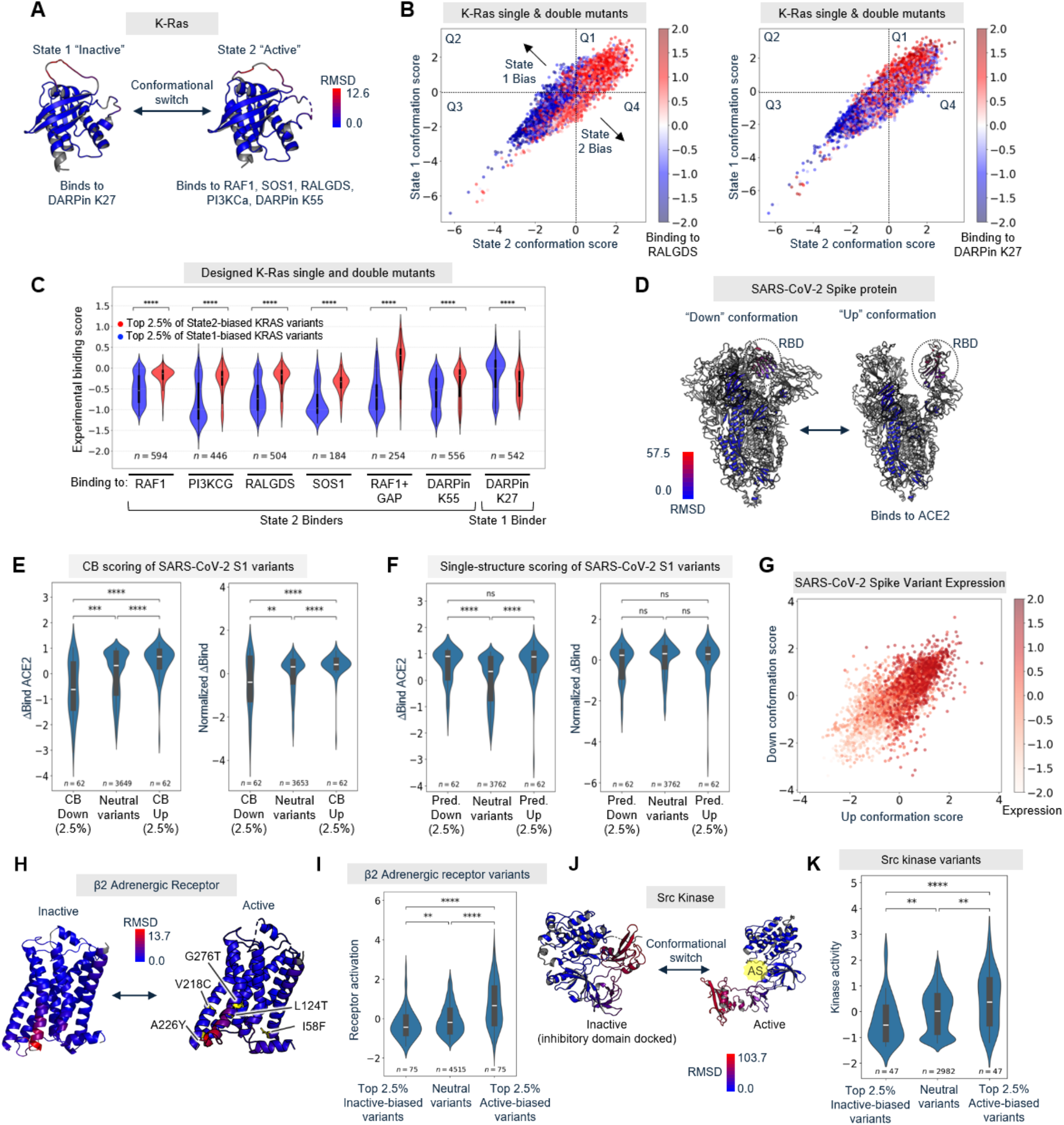
CB predicts protein variants with altered effector binding, receptor activation, and enzymatic activity. (**A**) The G protein K-Ras switches between an inactive State 1 conformation (PDB: 8T71, bound to GDP) and an active State 2 conformation (PDB: 6XHB, bound to GTP-analogue). State 2 binds to the effectors listed. RMSD, root mean square deviation. (**B**) 3320 single mutants and 26761 designed K-Ras double mutants were scored using CB against State 1 and State 2 experimental structures in (A). Points are colored according to experimental binding data from Weng et al.(24), for RALGDS, a State 2-specific effector (left) or DARPin K27, a State 1-specific binder (right). (**C**) Violin plot showing distributions of experimentally-determined binding scores from Weng et al.(24) for top CB-predicted State 1-biased variants (blue) and State 2-biased variants (red). Mutations at the KRAS-effector binding interface that are likely to directly influence effector binding are excluded from analysis (**Figure S1A**). ****p < 0.0001. Additional K-Ras analysis in Supporting Text and **Figure S1**. (**D**) SARS-CoV-2 spike protein with ACE2 receptor binding domain (RBD) in “down” (PDB: 7XIX) versus ACE2-binding competent “up” (PDB: 7XO8) conformations. (**E**) Distributions of raw ACE2 binding scores (left) and expression-normalized ACE2 binding scores (right), for CB-predicted up-biased and down-biased S1 variants. **p<0.01, ***p < 0.001, ****p<0.0001. CB scatter plot in **Figure S3A**. (**F**) Same as (E) except groups are based on single conformation scoring using ProteinMPNN instead of CB. ****p<0.0001. (**G**) SARS-CoV-2 RBD variants were scored using CB against RBD up/down structures. Points are colored by experimentally-determined variant expression levels from Starr et al.(25) (**H**) Human β2 adrenergic receptor (β2AR) in inactive and active conformations (PDB: 3SN6 and 2RH1, respectively). The conformational change in transmembrane helix 6 (red) enables the active form to bind to G proteins and arrestin. Positions of five CB-predicted active-biased mutations that are corroborated by experimental data from Jones et al (30) (Z > 1)are labeled. (**I**) 5263 β2AR variants were scored using CB, and top active-biased, inactive-biased, and neutral variants are plotted by their experimentally-determined receptor basal activation scores (30). ****p < 0.0001. (**J**) Src kinase repositions its auto-inhibitory SH2/SH3 domains upon autophosphorylation, becoming active (PDB: 2SRC and 1Y57, respectively). AS, active site. (**K**) Distributions of Src kinase activities for inactive-biased, active-biased, and neutral variant populations. **p < 0.01, ****p < 0.0001. CB scatter plot for Src in **Figure S3B**.

We next evaluated CB using a DMS dataset on SARS-CoV2 spike protein, whose receptor binding domain (RBD) binds to the mammalian ACE2 receptor in the “up” conformation but not “down” conformation (**Figure 2D**). Starr et al. evaluated 4022 point mutants of RBD, using yeast surface display to quantify their expression levels and ACE2 binding affinities(25). We used CB to calculate bias scores for all these variants, based on experimentally-determined structures of the S1 spike protein with RBD in the “up” or “down” conformation. We found that our CB-predicted top up-biased mutants have significantly (p<0.0001) higher ACE2 binding than a set of “neutral” variants predicted by CB to be neither up-biased nor down-biased (**Figure 2E**). Conversely, CB-predicted down-biased variants have reduced ACE2 binding scores compared to the neutral variant population.

Given the strong signal we observed without any additional modifications to the base model, we decided to compare CB’s performance to the use of IFMs as previously described, for scoring mutations on a single protein structure in the desired conformation. Such an approach led to the discovery of antibody variants with improved antigen affinity(18) and TEV protease variants with improved expression(19). We used ProteinMPNN to separately score RBD variants on the S1 “up” structure alone or “down” structure alone. When we analyzed the top scoring variants of these groups (same group sizes as those identified by CB) we found that their ACE binding scores were, surprisingly, both higher than that of the neutral population (**Figure 2F**, left). Since ProteinMPNN scores on both structures were highly correlated with variant expression level (**Figure 2G**), we repeated the analysis using ACE2 binding scores normalized by variant expression level. **Figure 2F** (right) shows that no significant differences were observed between groups in this analysis (for single-structure scoring), whereas the top up-biased and down-biased variants predicted by CB showed a significant increase and decrease in ACE2 binding, respectively, even when the binding scores were normalized by variant expression levels (**Figure 2E**, right). Thus, while conceptually simple, we believe the contrastive scoring objective used in CB is highly important for predicting conformational preference. We expand on this point in **Figure 5**.

We further validated CB across 6 additional DMS datasets, encompassing a G protein coupled receptor (GPCR)(26), a collection of 10 synthetic protein binder pairs(27), and four conformationally-gated enzymes(28-30). β2-adrenergic receptor (β2AR) undergoes a rearrangement of transmembrane helices upon activation that enables G protein binding, a characteristic shared by many other GPCRs (**Figure 2H**). We analyzed 5045 β2AR variants to predict mutations that bias the receptor towards its active versus inactive conformations, and found that our CB-predicted most active-biased and most inactive-biased variants showed significantly higher and lower basal receptor activities, respectively, compared to a neutral population (**Figure 2I**). **Figure 2H** shows that many CB-predicted biasing mutations with experimentally-validated effect on activation are distributed across the β2AR structure, and not just localized to TM6, which undergoes the largest conformational change upon receptor activation.

As a particularly challenging test of CB, we evaluated a collection of 10 synthetic protein binding pairs with highly similar sequences (hamming distances 1-8) and more importantly, and with only subtle changes in binding pose (aligned RMSD 0.23-2.6)(**Figure S2**)(27). CB was used to score the sequences of each binding pair on its own structure versus the conformations observed in the 9 other binding pair structures, to determine if a preference for the experimentally-validated true conformation could be detected. We found that most binder pairs were predicted to be biased towards their own experimentally-solved structure, with matched structure-sequence pairs scoring significantly (p < 0.01) higher than mismatched pairs. This difference was greater when structural/sequence differences were larger (e.g. L1.6/L1.1 vs. L2.17). Thus, we believe CB is sensitive to even very minor conformational changes and preferences.

Enzymes are dynamic machines that undergo structural rearrangements during catalysis to drive substrate binding, chemical bond formation or breakage, and product release. We identified four DMS datasets for enzymes that experience significant conformational change during regulation or catalysis: the kinases Src(30) and B-Raf(28), and the *E. coli* biosynthetic enzymes FabZ(29) and MurA(29). Src kinase switches between an auto-inhibited inactive conformation, mediated by phosphorylation of Tyr527, and an active conformation in which the auto-inhibitory domain is displaced (**Figure 2J**(31, 32). Indeed, we found that CB-predicted active-biased mutants have significantly higher activity than a neutral population, while inactive-biased mutants have significantly reduced activity; these effects are quantified in **Figure 2K**.

Inactive B-Raf, a kinase downstream of Ras, is monomeric and autoinhibited (33, 34). Upon activation by Ras, autoinhibition is released, resulting in B-Raf dimerization. Using CB on structures of B-Raf in both inactive monomeric and active dimeric states (**Figure S3C**)(34), we predicted dimer-biased and monomer-biased mutations. Although the DMS data from Simon et al.(28) is not complete and focuses primarily on mutations in the kinase domain of B-Raf, we found that predicted dimer-biased mutants showed a strong and significant increase in kinase activity (**Figure S3D**).

FabZ and MurA enzymes interconvert between open and closed structures, each associated with different stages of catalysis, and with minor structural differences between them (**Figures S3E, S3G**). Because a complete set of experimental structures was not available for these enzymes, we used AlphaFold-predicted templated structures for missing conformations (see Methods), which were just as effective for the implementation of CB. For both enzymes, the CB-predicted variants showed tuning of activity (as measured by bacterial growth) in the expected directions (**Figures S3F, H**). Collectively, our results suggest that CB can predict conformation-biasing mutations on diverse protein scaffolds, and that this effect can be harnessed to engineer or enhance protein activities.

### CB predicts conformationally-biased variants of lipoic acid ligase with highly altered labeling specificity

*E. coli* lipoic acid ligase (LplA) catalyzes sequence-specific lipoic acid attachment to acceptor proteins (**Figure 3A**), a process that is essential for bacterial metabolism and energy production(20, 21). We and others have previously exploited the malleability of LplA’s active site(35) to instead conjugate small molecule probes, such as fluorophores(22, 36) and Click chemistry handles(35, 37), to specific cellular proteins for live cell imaging, or to antibodies for therapeutic drug delivery (38). Crystal structures of LplA bound to lipoic acid(39), lipoyl-AMP ester(40), and H-protein acceptor(40) show that the enzyme undergoes a large-scale conformational change upon conversion of lipoic acid and ATP to lipoyl-AMP ester (adenylation step), that reconfigures the active site for the second step of catalysis, lipoyl transfer. A dramatic feature of this conformational change is the 180 degree rotation of the C-terminal domain (CTD), reorienting it from a downward, “closed” conformation that prevents docking of acceptor protein, to an upward “open” conformation that enables acceptor proteins to access bound lipoyl-AMP ester. We applied CB to LplA, both to experimentally evaluate CB’s ability to tune conformational occupancy, and to study the role of this conformational change in LplA’s catalytic activity and specificity.

**Figure 3.**
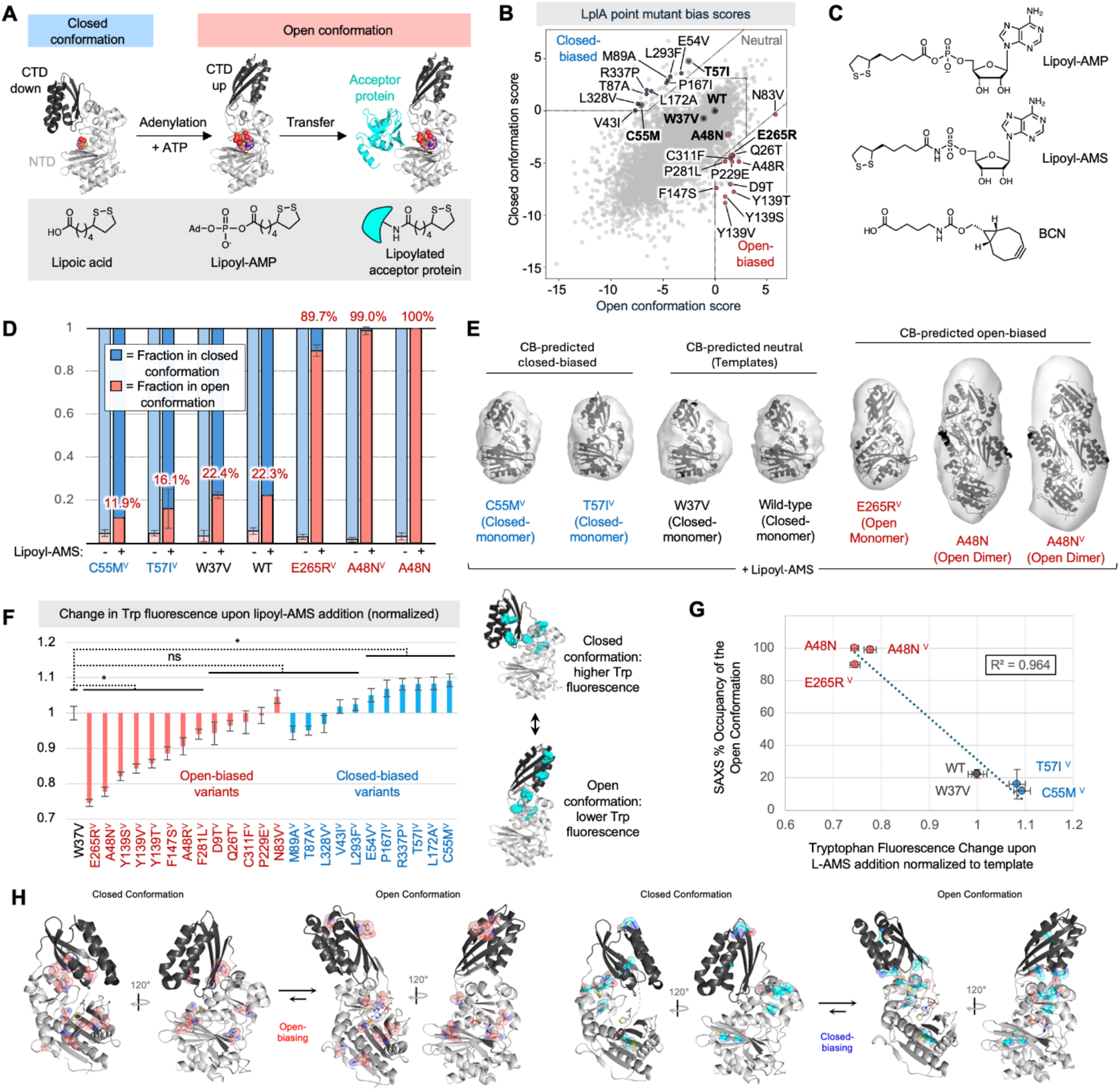
CB biases *E. coli* LplA towards open or closed adenylate ester-bound conformations. (**A**) Two half-reactions catalyzed by LplA: adenylation of lipoic acid by ATP to generate lipoyl-AMP, and transfer of lipoyl onto lysine sidechain of an acceptor protein. LplA switches from a closed to open conformation between the first and second half reactions. Ad, adenosine. CTD, C-terminal domain of LplA. NTD, N-terminal domain. Structures from PBD: 1×2H, 3A7R, and 3A7A (left to right). Lipoic acid, lipoyl-AMP, and octyl-AMP shown as red spheres (left to right). (**B**) CB on LplA, using open (3A7R) and closed (1×2G) backbone structures. Closed-biased, neutral, and open-biased variants selected for SAXS (bold) and Trp fluorescence analysis are labeled. (**C**) Structures of BCN, lipoyl-AMP, and non-hydrolyzable analog lipoyl-AMS. (**D**) SEC-SAXS analysis of four biased variants (on W37V background, indicated by V superscript), in apo state or in complex with lipoyl-AMS. Fractional occupancy predicted using Oligomer (see Methods and **Figure S5**), from 2-5 independent measurements per variant. Errors, ±1 std. dev. (**E**) DENSS *ab initio* modeling of protein envelopes, fit with major LplA conformer detected under +lipoyl-AMS condition. A48N appears as a dimer at the high protein concentrations required for SAXS (model generated by AlphaFold3). (**F**) Fold-change in tryptophan (Trp) fluorescence for purified LplA variants on addition of lipoyl-AMS. Open and closed structures of LplA show positions of Trp sidechains in light blue. n=3, p < 0.01. (**G**) Correlation of Trp fluorescence data from (F) to fractional occupancy predicted by SAXS Oligomer analysis in (D). (**H**) Distribution of top open-biasing (red) and closed-biasing (blue) mutations in LplA structure. **Figure S12** expands on the mechanisms by which these mutations may introduce conformational bias.

CB on LplA using open and closed crystal structures is shown in **Figure 3B**. When we applied CB to two different closed structures of LplA (in apo form or in complex with lipoic acid) as a control, the spread in the CB scatter plot was much reduced (**Figure S4A**). We also repeated the CB analysis using W37V LplA as a template instead of wild-type LplA, with largely identical results (**Figure S4B-D**). W37 is located at the end of LplA’s substrate binding pocket, and when mutated to valine, enables LplA to accept numerous unnatural probes in place of lipoic acid, including bicyclo-nonyne (BCN, **Figure 3C**)(41), a reactive handle for Click chemistry. CB-predicted mutations were introduced onto the W37V LplA template to allow for activity assays based on BCN ligation and Click derivatization by fluorophores or biotin.

We first established two assays for experimental evaluation of LplA’s conformational occupancy, which has not previously been characterized in solution. First, SEC-SAXS (size exclusion chromatography-small angle X-ray scattering), was used to measure LplA size and shape in solution, due to its ability to resolve conformational ensembles(42, 43) and suitability for mid-sized proteins like LplA (38 kD) that are too large for NMR(44) and too small for cryo-EM(45). Second, we developed a tryptophan (Trp) fluorescence assay to monitor LplA’s conformational change dynamically. This assay is also simpler to perform and requires less protein than SAXS.

We first characterized wild-type LplA and W37V LplA using both assays. SAXS scattering curves were fit with a modeled mixture of possible LplA conformers (**Figures 3D-E, S5**) (46), revealing that both enzymes predominantly adopt the closed conformation in the apo state and after lipoic acid binding - in agreement with published crystal structures(39). After complexation to lipoyl-AMP or its non-hydrolyzable analog, lipoyl-AMS (**Figure 3C**), however, both enzymes shifted only partially (22.3-22.4%) to the open conformation. Interestingly, this equilibrium is not reflected in the crystal structure, which shows LplA fully in the open conformation when bound to lipoyl-AMP. Generation of this crystal required seeding with LplA’s acceptor protein apoH(40), highlighting the non-physiological conditions of crystallography compared to solution measurements. Using the Trp fluorescence assay, we also observed the closed-to-open transition of wild-type and W37V LplA. **Figure S4F** shows a decrease in Trp fluorescence upon lipoyl-AMS addition, consistent with a shift towards the open conformation in which Trp sidechains become partially quenched by proximal arginine residues.

We then selected a representative set of 13 open-biasing mutations and 11 closed-biasing mutations predicted by CB (**Figure 3B**), and introduced them onto the W37V template (indicated by the V superscript) for recombinant expression, purification, and structural analysis. In our Trp fluorescence assay, most open-biased variants showed a greater drop in fluorescence upon lipoyl-AMS addition than W37V template, suggesting greater adoption of the open conformation (**Figure 3F**). On the other hand, most closed-biased variants changed less than W37V upon lipoyl-AMS addition, indicating a preference for remaining in the closed conformation. These results are highly consistent with the predictions of CB, with a correlation score of –0.61 (**Figure 5B**).

We then selected two strongly closed-biased variants and two strongly open-biased variants for further evaluation by SAXS (**Figure 4D-E**). Oligomer analysis indicated that all enzymes were predominantly in the closed conformation in the apo state, as expected. After incubation with lipoyl-AMS, the open-biased variants E265R^V^ and A48N^V^ shifted almost fully to the open conformation (90% and 99% open, respectively), a significant increase over wild-type and W37V LplA. Our closed-biased variants C55M^V^ and T57I^V^ showed the opposite trend, opening less than template W37V upon lipoyl-AMS addition, to just 12% and 16% open. Our SAXS data agreed with Trp measurements on the same LplA variants (**Figure 3G**). Together, our experimental characterization of conformational occupancy suggests that CB can effectively predict mutations that tune conformational equilibria.

**Figure 4.**
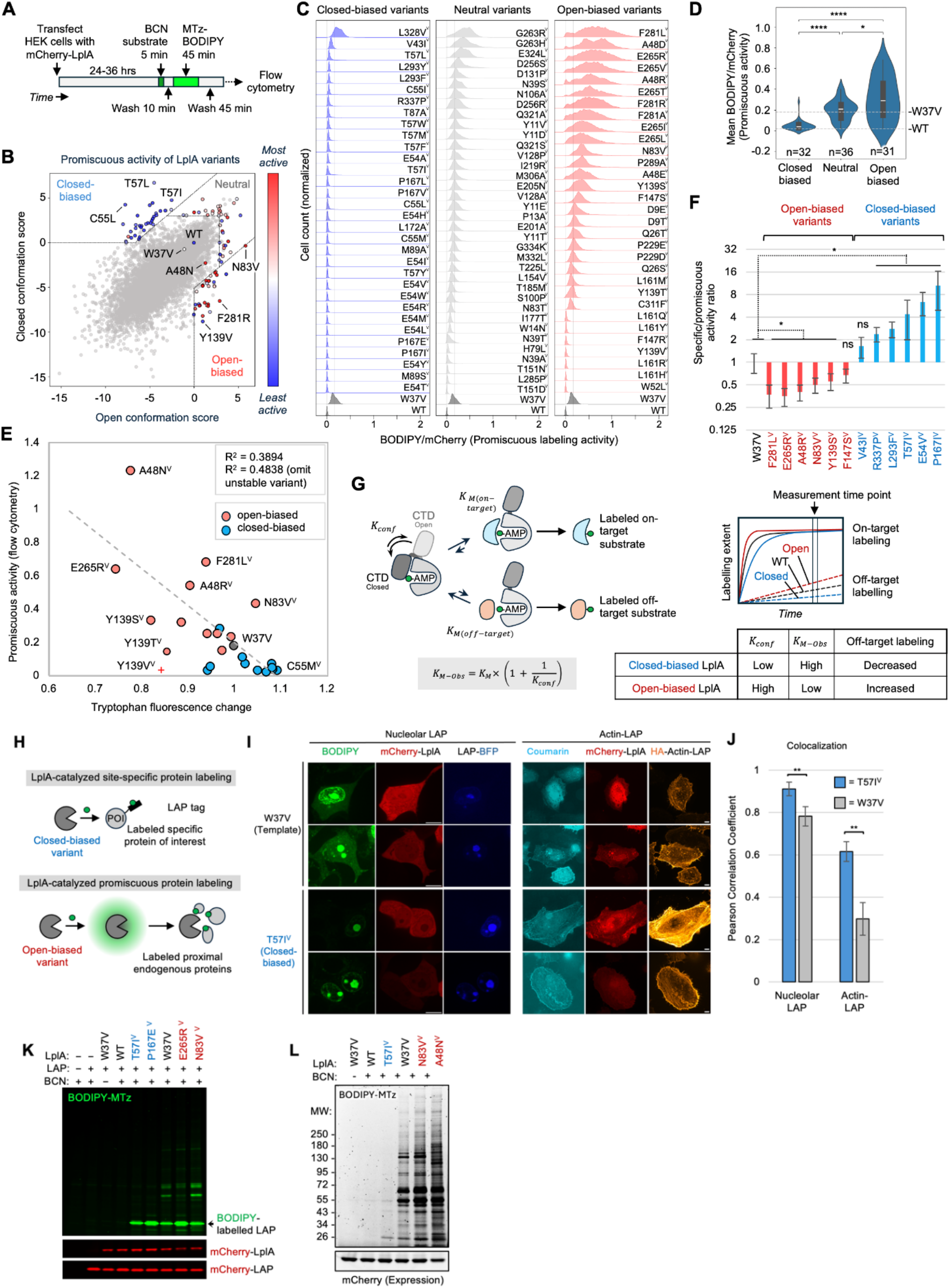
Conformationally-biased LplA variants show large differences in promiscuous protein labeling. (**A**) Flow cytometry assay for measuring the promiscuous labeling activity of mCherry-tagged LplA variants. After conjugation to endogenous proteins, BCN is derivatized with fluorogenic methyltetrazine(MTz)-BODIPY. See **Supplementary Text** and **Figure S7** for further discussion of this assay. (**B**) CB plot for LplA, with closed-biased, neutral, and open-biased mutants colored by their experimentally-measured promiscuous activities in (C). (**C**) Promiscuous activity histograms for closed, neutral, and open-biased LplA variants (all variants made on W37V background except WT). 2D BODIPY vs. mCherry flow cytometry plots for all variants shown in **Figure S8**. This experiment was performed 2 times with similar results (correlation between replicates shown in **Figure S6D**). (**D**) Mean promiscuous activity scores for each category of CB-designed LplA variant in (C). * p<0.05, **** p<0.0001. (**E**) Correlation between promiscuous activity from (C) and Trp fluorescence change measured in **Figure 3F**. Y139V (red cross) is an unstable variant that shows strong aggregation in HEK293T cells. (**F**) Ratio of specific to promiscuous activity for 12 purified LplA variants, based on in-cell promiscuous activity measurements in (C) and E2p specific labeling assay in **Figure S6F-G**. Errors, 1 std. dev. *p < 0.05. (**G**) Proposed mechanistic model for how conformational occupancy (*K*_*conf*_) tunes substrate specificity in LplA. In lipoyl-AMP-bound LplA, the C-terminal domain (CTD) may act as a tethered competitive inhibitor, increasing *K*_*M-Obs*_ for protein substrates when *K*_*conf*_ (equilibrium constant between closed and open conformations) is low. Modeling of on-target and off-target labeling velocities shown in **Figures S6J-K**. (**H**) LplA variants can be used for either site-specific (top) or promiscuous (bottom) protein labeling in cells. POI, protein of interest. LAP is a 13-amino acid engineered LplA Acceptor Peptide(47). (**I**) Confocal microscopy of LAP fusion proteins tagged with coumarin or BODIPY fluorophores, catalyzed by W37V LplA (top) or closed-biased variant T57I^V^ (bottom). For BODIPY labeling (left), HEK293T cells were treated with BCN for 10 minutes, then clicked with BODIPY-MTz, and imaged live. For coumarin (right), HeLa cells were labeled for 10 minutes with coumarin-AM2(22), then fixed and stained with anti-HA antibody to visualize actin-LAP. Scale bars, 10 um. (**J**) Quantification of images in (I), analyzing colocalization between BODIPY and BFP (left) or coumarin and HA (right). 4 FOV per condition. ** p < 0.005. (**K**) In-gel fluorescence of cell lysates labeled as in (I). (**L**) The open-biased LplA variants A48N and N83V catalyze promiscuous labeling of endogenous cytosolic proteins in HEK293T cells. Cells were treated with BCN for 10 minutes before lysis, Click with mTz-BODIPY, and SDS-PAGE. No LAP fusion was expressed in these samples.

Next we explored the functional consequences of biasing LplA’s conformation. Because the C-terminal domain (CTD) of LplA blocks acceptor protein access in the closed but not open conformation, we hypothesized that open-biased variants might be either more active than WT, or able to tag a wider range of proteins (i.e., more promiscuous). To test this, we developed a flow cytometry-based assay to evaluate the promiscuous labeling activities of LplA variants in the cytosol of living mammalian cells. As shown in **Figure 4A**, mCherry-tagged LplA variants are expressed in HEK293T cells, and membrane-permeable BCN substrate is supplied for 5 minutes (the cell provides ATP). If LplA conjugates BCN to endogenous mammalian proteins, we can detect such labeling through subsequent Click reaction with the BCN-reactive methyltetrazine(MTz)-BODIPY fluorophore. Labeled cells are quantified by flow cytometry and expression-normalized promiscuous labeling activities (BODIPY/mCherry ratio) are plotted as a histogram.

From the CB plot in **Figure 4B**, we selected the top 31 open-biasing, 32 closed-biasing, and 36 neutral variants (high in both open and closed scores) for experimental evaluation on a W37V background. This set includes the variants we tested above by SAXS and Trp fluorescence. The flow cytometry histograms in **Figure 4C** show that the majority of open-biased variants exhibit increased promiscuous labeling compared to the template W37V, while nearly all closed-biased variants show reduced promiscuous activity. For several open-biased variants with low activity, structural analysis suggests that they may have steric clashes with BCN substrate (**Figure S6I**). Neutral variants had promiscuous activities in between those of the open-biased and closed-biased variants (**Figure 4D**).

We also evaluated the *specific* labeling activities of a subset of these LplA variants. 13 of the open- and closed-biased variants that we analyzed by Trp fluorescence were expressed in HEK293T cells along with E2p, an acceptor protein domain from one of LplA’s endogenous bacterial substrates, pyruvate dehydrogenase(21). After 5 minute labeling with BCN, cells were lysed, Clicked with mTz-BODIPY, and analyzed by SDS-PAGE. **Figure S6F-G** shows that all variants labeled E2p to a similar extent as W37V template. Thus, the target-specific labeling activities of these variants are not measurably changed, in contrast to their dramatically altered promiscuous labeling activities (**Figure 4F**). The good correlation between measured promiscuous activity and Trp fluorescence change across LplA variants (**Figure 4E**) suggests that LplA’s sequence-specificity is closely linked to changes in its conformational equilibria.

A possible mechanistic model is presented in **Figure 4G**. The C-terminal domain of LplA is required for its adenylation activity (20), but after generation of lipoyl-AMP ester, its position over the active site in the closed conformation inhibits acceptor protein access for the second transfer step.

Thus, the CTD can be modeled as a tethered competitive inhibitor of lipoyl transfer. As such, its effect is to increase the effective *K*_*M-Obs*_ of LplA for its protein substrates by the equation shown(48). Modeling in **Figures S6J-K** shows that under the assumption that *K*_*M*_ values for on-target substrates are low (*K*_*M on-target*_ << [S_on-target_]) and *K*_*M*_ values for off-target substrates are high (*K*_*M off-target*_ >> [S_off-target_]), the ratio of their initial rates (V_on-target_/V_off-target_) will change with *K*_*Conf*_, the equilibrium constant for the closed to open conformational transition. Closed-biased variants have lower *K*_*Conf*_ (are more inhibited by the CTD), higher effective *K*_*M-Obs*_, and thus label off-target proteins less than wild-type or W37V LplA. By contrast, open-biased LplA variants have less inhibition or no inhibition, lower effective *K*_*M-Obs*_, and can readily transfer lipoic acid or BCN to a wide range of off-target substrates. The net result is more promiscuous labeling by open-biased variants, and more specific labeling by closed-biased variants of LplA.

Because W37V LplA has been widely used as a biotechnology for site-specific protein labeling with small-molecule probes in cells(22, 35-37) and in vitro(38, 49) (**Figure 4H**), the changes in substrate specificity introduced by CB could be highly useful for either decreasing or increasing off-target protein labeling in such applications. We explored this by first using LplA to specifically conjugate fluorophores onto LAP fusion proteins in mammalian cells. LAP is an engineered 13-amino acid peptide that is recognized by LplA (47)and has been used to target coumarin(22), resorufin(36), Click handles(35, 37) and photocrosslinkers(50) onto proteins of interest. When we performed in-cell labeling of membrane-targeted LAP-BFP-CAAX (CAAX is a lipidation sequence) with W37V LplA and BCN, followed by mTz-BODIPY Click, we observed considerable background due to the slight promiscuous activity of W37V, especially at high expression levels (**Figure 4I-J**). When we replaced W37V with the CB-predicted closed-biased variant T57I, however, the signal to noise (S/N) of labeling was significantly improved (p < 0.005), consistent with the increased specificity of T57I. The difference was also apparent from SDS-PAGE analysis of labeled cell lysates, showing a single labeled band for T57I (corresponding to the LAP fusion protein) but labeling of multiple bands for W37V and open-biased variants (**Figure 4K**). Thus, closed-biased LplA variants can usefully improve signal-to-noise in applications requiring high sequence-specificity.

On the other hand, promiscuous open-biased LplA variants could be useful for a different protein labeling technology: proximity labeling (PL)(51). Promiscuous enzymes such as TurboID(52) and APEX(53) have been widely applied for PL-based mapping of organelle proteomes and protein interactomes in living cells and animals. In separate work (unpublished), we engineered the new PL enzyme “FlexID” from open-biased variants of LplA. FlexID overcomes some of the important limitations of existing PL technologies, such as the high biotin background associated with TurboID(51, 52), the toxic labeling conditions of APEX, and the slow speed of BioID. **Figures 4L, S6C** show how open-biased LplA variants can catalyze the labeling of hundreds of endogenous proteins in the cell compartments in which they are expressed. Overall, our application of CB to LplA shows that: (1) the conformational equilibria of complex, multistep enzymes can be tuned; (2) conformational biasing can produce dramatic changes in enzyme function; and (3) biased enzyme variants can be harnessed for useful biotechnological applications.

### Benchmarking CB

To assess the quality of CB’s predictions in the context of LplA, we plotted CB bias scores against three experimental datasets: SAXS data for 6 LplA variants (**Figure 5A**), Trp fluorescence measurements for 24 variants (**Figure 5B**), and mean promiscuous labeling activities for 99 variants (**Figure 5C**). All three datasets show good correlation with CB bias score. For the 31 open-biased variants we evaluated by flow cytometry, 25 should be free from steric clashes with BCN substrate. Of these, 23/25 (over 90%) show either increased promiscuity or increased tryptophan fluorescence change, consistent with open bias. Likewise, 31out of 32 closed-biased variants (over 95%) show the expected decrease in promiscuity. Thus, variants with the most divergent CB scores are highly enriched for the intended conformation-specific functions.

**Figure 5.**
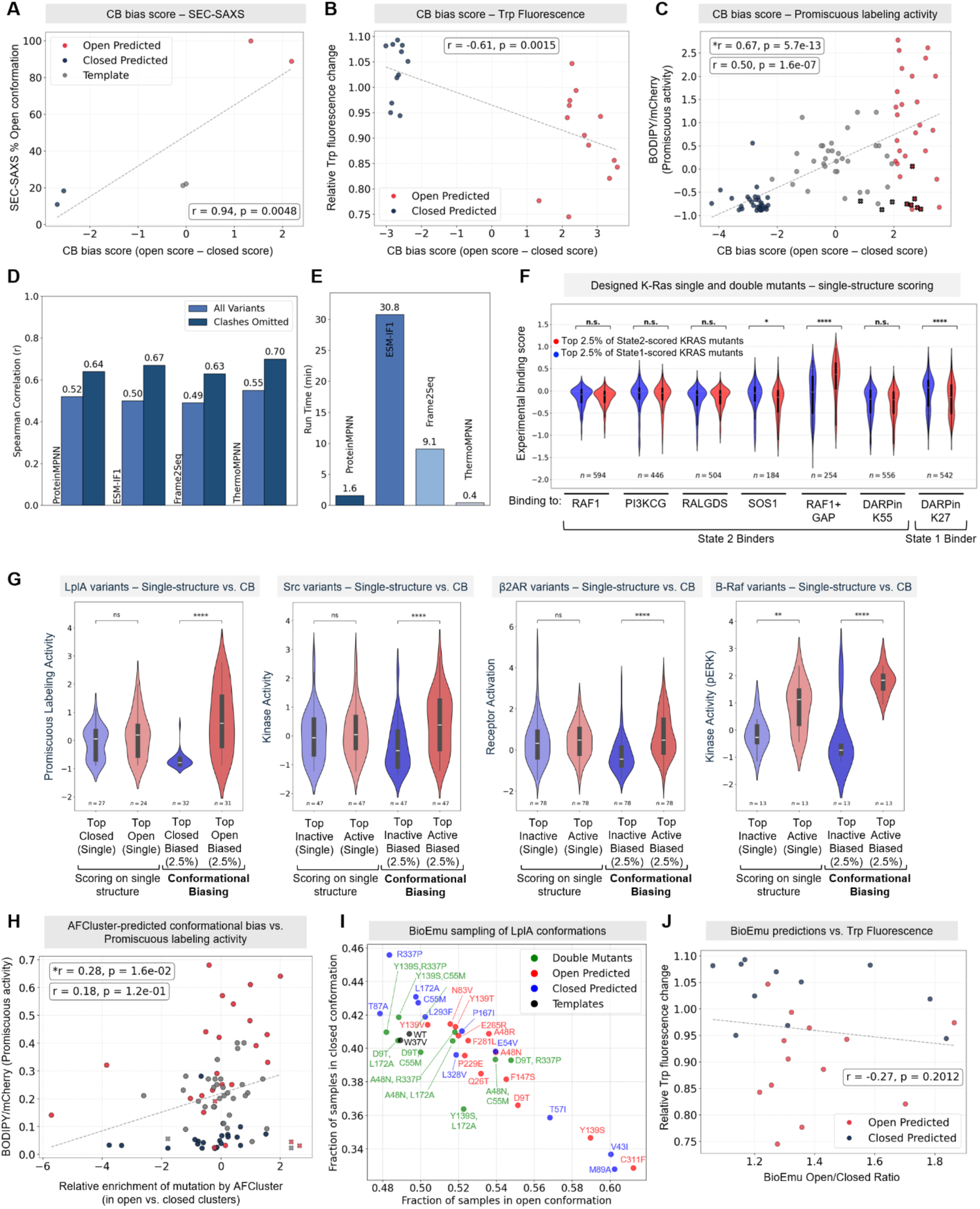
Benchmarking CB. CB bias scores for LplA plotted against experimental data from (**A**) SEC-SAXS, (**B**) Trp fluorescence, and (**C**) promiscuous labeling activity measurements. Variants are colored by CB-prediction: closed-biased (blue), neutral (grey), and open-biased (red). Variants marked with an X are likely to have a steric clash with BCN substrate in the active site (AS), explaining their low activities (**Figure S6I**). *r = spearman correlation calculated without steric clash variants, r = spearman correlation calculated for all variants. (**D**) Correlation between LplA promiscuous activities and CB scores, determined using ProteinMPNN, ESM-IF1, Frame2Seq, or ThermoMPNN. (**E**) CB runtime on a single RTX 4090 GPU using different inverse folding models, in minutes. Data was processed using consistent parameters. (**F**) K-Ras analysis performed using single structure scoring. *p<0.05, ****p<0.0001. Compare to **Figure 2C**. (**G**) Comparison of CB vs. single structure scoring using ProteinMPNN, for LplA, β2AR, Src, and Braf datasets. **p<0.01, ****p<0.0001. (**H**) Correlation between AFCluster-determined bias scores (difference in open vs. closed structural similarity score) and LplA promiscuous labeling activities. (**I**) BioEmu sampling (n=500) of 12 CB-predicted open-biased variants (red) and 12 CB-predicted closed-biased variants (blue). Samples were aligned to LplA open and closed structures to estimate fractional open/closed occupancy, which is plotted. Green points are combination mutants, in which a closed-biasing mutation is combined with an open-biasing mutation in the same protein. These were simulated (n=100) using BioEmu and plotted in the same manner. (**J**) Correlation between BioEmu predictions and LplA Trp fluorescence data for 24 variants. Red points are predicted by CB to be open-biased, and blue points are predicted to be closed-biased. For comparison, CB vs. Trp fluorescence correlation plot is in (B).

Though we selected ProteinMPNN as our IFM for CB scoring, CB is in principle agnostic as to the specific IFM used. To test if our predictions were generalizable across IFMs, we evaluated three other IFMs for CB scoring in place of ProteinMPNN. We selected two masked language models, ESM-IF1(16) and Frame2Seq(54), as well as ThermoMPNN(55), a version of ProteinMPNN fine-tuned on thermal stability data. Since ThermoMPNN is specifically designed for predicting ΔΔG, it may be especially sensitive to the state-specific stabilizing effects of specific variants. We used both LplA and β2AR datasets, and compared the agreement of CB bias scores with experimental data. **Figure 5D** shows that for LplA, all IFMs give good correlation with LplA mean promiscuous activities (spearman correlations between 0.63-0.7, excluding substrate clash variants; full scatter plots in **Figure S9A-D**). For β2AR, ProteinMPNN and ThermoMPNN gave the most significant increase in active-biased predicted mutant activity, while Frame2Seq gave predictions that deviated the furthest from the other models (**Figure S9E-G**). All models were able to enrich for active-biased mutations. Interestingly, the 18 active-biased variants predicted by all four models had even higher receptor activation scores than the original CB prediction group based on ProteinMPNN scoring alone (**Figure S9H**).

While no gold standard method for predicting conformation-biasing mutations has been established, we attempted to benchmark CB using three alternative methods for conformational scoring: single structure IFM scoring, AFCluster(8), and BioEmu(56). Single structure IFM scoring was already applied to SARS-CoV2 variants in **Figure 2F**. We extended our analyses to LplA, K-Ras(24), B-Raf(28), and β2AR(26)datasets as well (**Figure 5G**). For LplA, we used ProteinMPNN to predict highest-likelihood mutations in the wild-type sequence, using either the open structure alone or the closed structure alone. 24/31 top-scoring variants on the open structure and 27/32 top-scoring variants on the closed structure were present in our promiscuous labeling dataset. **Figure 5G** shows that these groups do not exhibit a significant difference in promiscuous labeling activities. On the other hand, CB’s contrastive scoring method was able to predict variants with increased and decreased promiscuous labeling activities (for open- and closed-biased variants, respectively). The same trends were observed for K-Ras, B-Raf, and β2AR (**Figure 5G**), demonstrating the importance of CB’s contrastive scoring compared to traditional single structure IFM scoring for predicting conformation-specific effects.

We next assessed AFCluster(8) for the prediction of conformationally-biased variants of LplA. First, we used AFCluster to generate clusters of similar sequences from LplA’s multiple sequence alignment (MSA). We then scored each cluster’s AlphaFold2-predicted structure for similarity to LplA’s open or closed conformations (**Figure S10**). By quantifying the enrichment of individual mutations in open-or closed-like clusters as described in the AFCluster study, we estimated conformational bias scores (see Methods). We found that these correlated much more poorly with our experimental data than CB bias scores did (**Figure 5H**, r = 0.26 compared to r = 0.67 for CB in **Figure 5C**). Furthermore, 1250 LplA point mutants (20%) could not be scored by AFCluster at all because they are absent from LplA’s MSA; this list includes many biasing mutations predicted by CB and validated experimentally. Our analysis highlights a key limitation of AFCluster – that it requires evolutionary branches in which the protein of interest adopts alternate ground-state conformations, a feature not present in many conformation-changing proteins, including LplA.

Next, we compared CB to BioEmu(56), a state-of-the-art model for sampling equilibrium conformational ensembles of proteins. We used BioEmu to generate 500 structural samples for each of the 24 LplA variant sequences that we analyzed by Trp fluorescence in **Figure 3F**. By analyzing the conformational distributions of these samples, we estimated a bias score for each variant by taking the ratio of the number of samples generated in the open conformation to the number generated in the closed conformation (**Figure 5I, S11A**). While we observed some agreement between BioEmu’s bias scores and our Trp fluorescence data (spearman correlation -0.27, **Figure 5J**), the correlation was worse than that observed using CB bias scores (spearman correlation -0.61, **Figure 5B**). BioEmu is also orders of magnitude more computationally intensive, making it impractical to analyze thousands of protein variants, and it requires additional interpretation of sampled structures.

BioEmu is trained to sample diverse conformational states for a given protein. We observed from our BioEmu analysis of LplA that for all 24 variants analyzed, the vast majority (∼90%) of sampled structures occupied the same set of native conformations as wild-type LplA, appearing as open-like, closed-like, or in transition between open and closed states, with <5% deviation between variants (**Figure S11B**). We wondered how BioEmu predictions would look for double mutants combining open-biasing mutations with closed-biasing mutations in the same LplA sequence. Sampling (n=100) of double mutants gave predictions of conformational occupancy in between those of the original mutations, supporting the idea that CB-predicted mutations act on the same conformational equilibrium in opposing directions (**Figure 5I**).

We further applied BioEmu to analyze CB-predicted SARS-CoV2 RBD variants (25, 57, 58). We selected a high-confidence subset of CB-predicted “up”-biased and “down”-biased RBD variants with increased and decreased ACE binding, respectively, avoiding mutations in the binding interface with ACE2 (**Figure S11C**). Using BioEmu, we generated an ensemble (n=4000) of conformational states for each variant, and classified them by structural topology as “up”-like or “down”-like. We observed that for these high-confidence variants predicted by CB, BioEmu predictions were largely in agreement, and showed good correlation with the experimental data (r = 0.82, **Figure S11D**), supporting our previous predictions.

Overall, we believe that CB shows a strong ability to predict conformationally biasing mutations compared to other alternatives, while reflecting a potentially generalizable and important application of IFMs. In addition, CB is significantly more efficient in terms of computational requirements, rendering it immediately applicable to a wide range of biological problems.

## Discussion

In this study, we developed Conformational Biasing (CB), a computational method to identify mutations that bias the conformational equilibria of proteins to produce improved or altered functions. We validated CB on 7 different deep mutational scanning (DMS) datasets by predicting variants that increase conformation-specific functionality, such as effector binding for K-Ras in its State 2 conformation, reporter gene expression downstream of the β2 adrenergic receptor in its active signaling conformation, and enzymatic activity of Src kinase in its uninhibited conformation. In addition, we showed that CB could differentiate between 10 highly similar synthetic protein binding pairs. Finally, we applied CB to modulate the conformational equilibria of lipoic acid ligase in its adenylate ester-bound intermediate state, and discovered that doing so dramatically altered its on-target/off-target labeling ratio. Higher specificity “closed”-conformation biased variants were used for more specific fluorophore tagging of LAP fusion proteins in living cells, while more promiscuous “open”-conformation biased variants were used to develop proximity labeling technology for spatial proteome and interactome mapping. In addition to uncovering a previously unknown mechanism for conformational gating of LplA’s specificity, CB produced useful reagents for biotechnology.

For the development of CB, we prioritized scalability, accessibility, and generalizability, which partially motivated our initial interest in inverse folding models. CB can analyze >3000 variants in under a minute on a consumer-grade GPU (RTX 4090), and is easy to run using free resources like Google Colab. We have created a Colab notebook to run the CB workflow (https://github.com/alicetinglab/ConformationalBiasing/) in order to maximize accessibility.

We were encouraged to find that CB produced signal on every dataset that we tested, despite vast differences in protein functions (G protein, virus, receptor, kinase, ligase, binder), protein size, oligomeric state, and magnitude of the conformational change. Structures used for CB scoring could be either experimental structures or AlphaFold-predicted structures, as long as they capture conformational states with functional relevance. For all but our LplA dataset, the readouts of conformational equilibria were indirect – via effector binding or receptor activation linked to specific conformational states, for example – which is an added layer of noise; yet CB was still able to enrich for biasing mutations. Since conformational change is tied intimately to so many different protein functions, CB has the potential to improve or alter many diverse protein types.

We were surprised to find that single point mutations identified by CB could so dramatically alter conformational equilibria, sometimes with large functional consequences. We examined the location and nature of CB-predicted mutations in LplA (**Figures 3H** and **S12**). Some mutations appear to favor the open conformation of LplA by introducing steric collisions (N83V) or repulsive interactions (E265R) that disfavor the closed conformation. Other mutations, such as open-biasing P289A, cause a loss of beneficial hydrophobic packing in the closed state. For other CB-predicted mutations, however, the mechanistic basis of biasing is unclear. Many CB mutations are far from the region that undergoes the largest conformational change (for example, in **Figures 2H** and **3H**), but experimental data validates that they are effective. These variants may tap into long-range allosteric networks that are not yet appreciated in the proteins under study. As such, CB has the potential to be a tool for discovery of hotspots and mechanisms of conformational switching in proteins.

Inverse folding models like ProteinMPNN have been widely used for the de novo design of sequences that fold into desired backbone structures(15, 59). However, the role of IFMs in the design and evaluation of targeted mutations may be just as important. For example, IFMs have been applied to antibody-antigen complexes for unsupervised prediction of higher affinity mutants(30). CB differs from this interesting use case in two key ways. First, instead of using IFMs to improve the fitness of a sequence, which is often highly related to stability and expression, CB takes a more targeted approach to modify a specific property (conformational equilibria), based on the hypothesis that conformational occupancy is tied to function such as enzymatic activity or specificity. Second, we showed that CB’s contrastive scoring is much more effective than single structure scoring by IFMs for predicting variants with altered functionality. For example, when ProteinMPNN was used to separately predict the best mutations on the active conformation or inactive conformation of proteins of interest, the resulting groups did not, in most cases, show higher or lower functional activities; instead, they often showed just improved expression (**Figures 2F-G, 5G**). We found that mutations in LplA that scored highly on both open and closed conformations were not as active for promiscuous labeling as open-biased mutations predicted by CB. This suggests that, at least in some cases, designing conformational occupancy rather than absolute fitness may be more appropriate to achieve design goals.

A growing number of computational tools are being developed that consider more than one conformation of a protein of interest. We benchmarked CB using two such state-of-the-art tools, AFCluster(8) and BioEmu(56). AFCluster predictions had minimal signal when compared to our LplA experimental data, and the methodology is limited by its strong reliance on MSA sub-clustering. By contrast, CB can predict biasing mutations outside of the evolutionary record. BioEmu was able to recapitulate some of CB’s predictions on our LplA experimental data, but the correlation of BioEmu scores with experimental measurements was still poorer than that of CB scores. With comparable computational resources (such as those accessible through Google Colab), CB can be run on a model system on the timescale of minutes, while a comparable analysis with BioEmu could take multiple days.

We found that the signal from CB was not specific to a single IFM, but rather seems to be a task that IFMs are generally suited to. While CB currently has multiple limitations, such as the requirement for high-quality experimental structures or predicted structures in multiple states, and limited predictive power throughout all point mutants, we anticipate that these will improve as both IFMs and structure prediction methods improve. Extensions of our CB method, whether with a next-generation IFM, more complex training data, inclusion of side-chain and ligand modelling, or integration of protein language models, could further improve performance and unlock new applications.

## Acknowledgements

We are grateful for funding from the National Institutes of Health (RC2DK129964), the Chan Zuckerberg Biohub – San Francisco, the GPCR collaborative of the St. Jude Children’s Research Hospital, the Phil and Penny Knight Initiative for Brain Resilience, the Wu Tsai Neurosciences Institute, and Stanford Bio-X. P.E.C. is supported by an NSF Graduate Research Fellowship. Dan Herschlag, Brian Trippe, Brian Hie, James Ferrell, Chris Garcia, Matthew Ravalin, Jessica Simon, Zhaoyang Li, and Nicholas Kalogriopoulos provided valuable feedback. We thank Reika Tei for assistance with LplA microscopy, Pratik Kumar and Luke Lavis for sharing JF-646 tetrazine fluorophore.

## Conflict of interest statement

The authors declare no conflict of interest.

## Author contributions

P.E.C., A.G.X., and A.Y.T. conceived of the project and developed the CB methodology. A.G.X. performed CB predictions and analyses. A.G.X. and P.E.C analyzed deep mutational scan datasets, with input from A.Y.T. and S.D.. A.G.X. performed CB benchmarking. P.E.C., A.G.X., and A.Y.T. designed the LplA library. P.E.C. and A.Q. designed and executed the cell-based characterization of LplA variants. S.D. developed the Trp fluorescence assay and LplA purification method, and synthesized coumarin and lipoyl-AMS. P.E.C. purified LplA variants and performed the Trp fluorescence assay. P.E.C. and T.M. performed SEC-SAXS. P.E.C., T.M., and A.G.X. analyzed SAXS data. P.E.C., A.G.X., and A.Y.T. wrote the manuscript with input from all authors.

## Supplementary Materials

Figs. S1 to S12

Supplementary Tables 1-9 (as .xlsx file)

Supplementary Text

Materials and Methods

## Notes

### Competing Interest Statement

The authors have declared no competing interest.

### Summary of Updates

This version of the manuscript has been revised to include new data and analyses.

