## Supplementary Information for "Computational design of conformation-biasing mutations to alter protein functions"

### Supplementary Material for
### Computational design of conformation-biasing mutations to alter protein functions

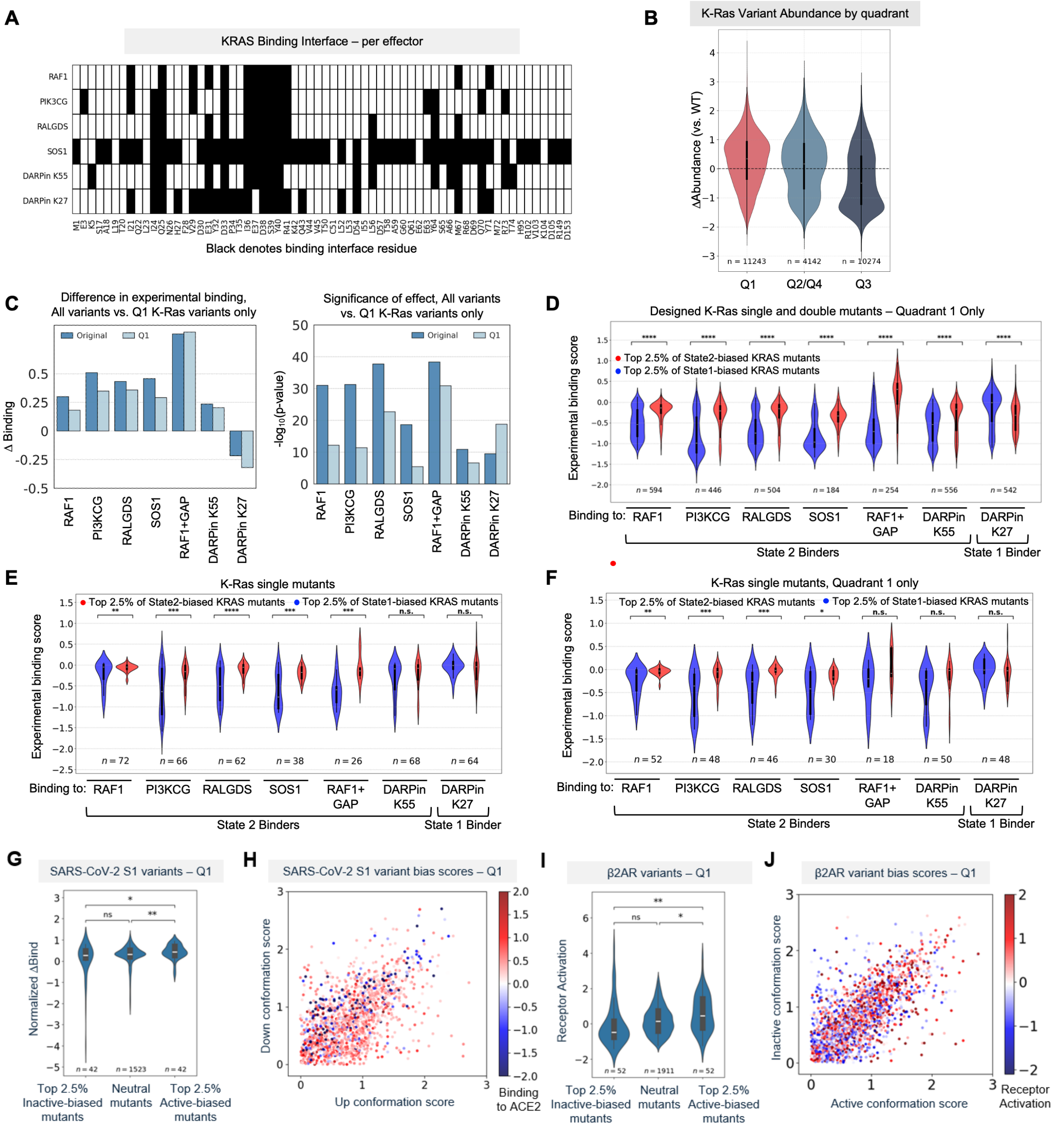

**Figure S1. CB analysis of Q1 variants of K-Ras, SARS-CoV2 spike protein, and β2AR.** (**A**) Residues shown in black represent previously-determined binding interface residues as reported in Weng et al. (1), or residues directly adjacent to two binding interface residues (high likelihood of interaction with effector). These sites are removed in the analysis of K-Ras variants (see Methods). (**B**) Experimentally-measured (1) K-Ras variant abundance, segregated by score on State 1/State 2 structures (quadrants labeled in **Figure 2B**). (**C**) Change in experimental binding scores and $Log_{10}$ P-values when comparing effector binding scores for top State 1-biased vs. State 2-biased K-Ras variants, for all variants (dark blue) or for Quadrant 1 variants only (light blue). (**D**) Same as **Figure 2C**, but only Quadrant 1 or Q1 (variants with above-average score on both conformations) variants are analyzed ****p<0.0001. (**E**) Same as **Figure 2C**, but only single mutants are analyzed **p<0.01, ***p<0.001, ****p<0.0001. (**F**) Same as (D) but analyzing Q1 single mutants only *p<0.05, **p<0.01, ***p<0.001 (**G**) Comparison of ACE2 binding scores for the CB-predicted most-biased SARS-CoV2 variants in Q1 only. *p<0.05, **p<0.01 (**H**) Scatter plot of Q1 variants, colored by expression-normalized ACE2 binding scores from Starr et al.(2). (**I**) Comparison of β2AR receptor activation scores for CB-predicted most-biased variants in Q1. *p<0.05, **p<0.01 (**J**) Scatter plot of Q1 β2AR variants, colored by experimentally-measured receptor activation(3).

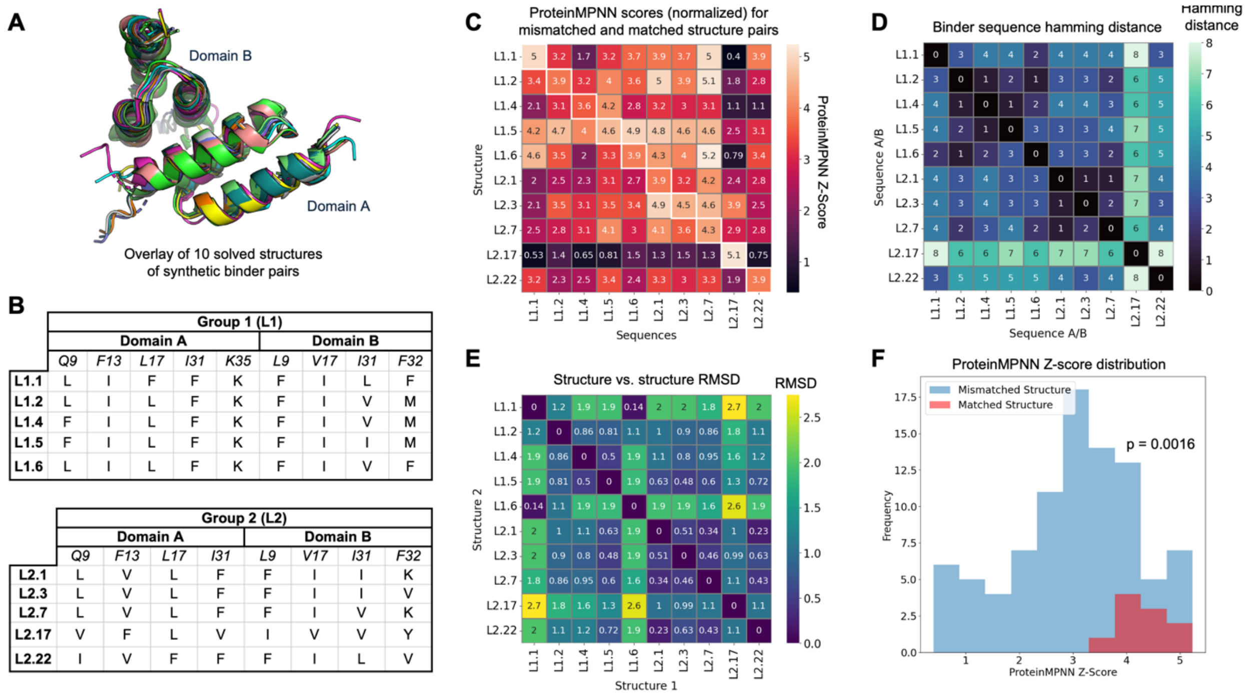

**Figure S2. CB predicts the conformational preferences of synthetic protein binder pairs.** (**A**) Overlaid structures of 10 synthetic binder pairs from Yang et al., derived from Staphylococcus Protein Z and its designed affibody binder(4). (**B**) Table summarizing mutational differences between binder pairs. Domain A binds to Domain B. (**C**) Heatmap showing results of CB, scoring each binder pair sequence against its corresponding structure or a mismatched structure. Scores were scaled and normalized relative to a synthetic distribution of generated sequences (see Methods). (**D**) Hamming distances between binder pairs, showing high sequence similarity. (**E**) RMSD of aligned structures. (**F**) Histogram showing the distribution of CB-generated ProteinMPNN scores for matched versus mismatched sequence-structure pairs. Significance of difference between distributions evaluated by Student's t-test.

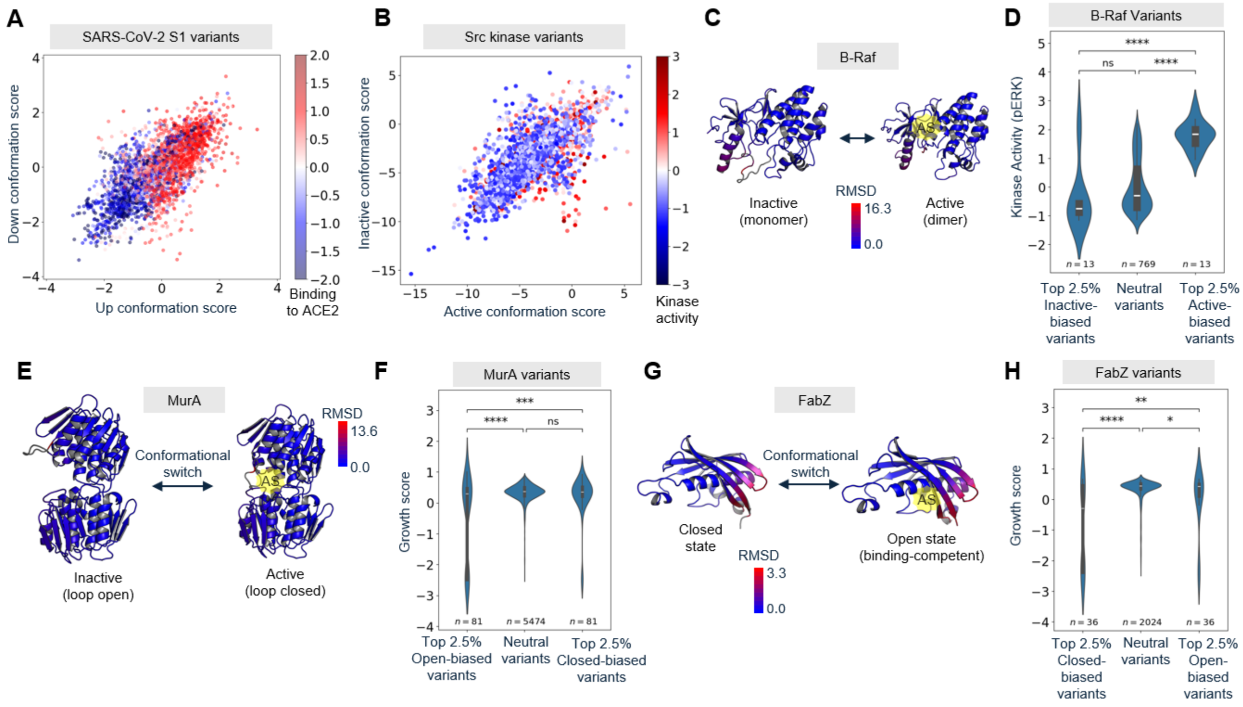

**Figure S3. Additional data related to Figure 2.** (**A**) SARS-CoV-2 RBD variants were scored using CB against RBD up/down structures in **Figure 2E**. Points are colored by experimentally-determined ACE2 binding scores from Starr et al.(2) (**B**) 3372 Src kinase variants were scored using CB and the inactive and active structures shown in **Figure 2J**. Points are colored according to experimentally-determined kinase activity data (based on inhibition of cell growth) from Ahler et al.(5). (**C**) B-Raf is a human kinase downstream of Ras, which has an inactive (autoinhibited, monomeric) state, and an active (phosphorylated, dimerized) state (PDB: 7MFD and 7MFF, respectively). AS, active site. (**D**) Distributions of experimentally-determined B-Raf activities from Simon et al.(6), based on quantification of phospho-ERK levels. Categories are based on CB analysis of active and inactive B-Raf structures in (D). ****p < 0.0001. (**E**) MurA is an *E. coli* transferase required for peptidoglycan synthesis that interconverts between an inactive "loop open" state and an active "loop closed" conformation in which substrates are re-positioned for catalysis (PBD: 1UAE). The "loop open" structure was generated by AlphaFold2, templating on the open structure of *E. cloacae* MurA (PBD: 1NAW). AS, active site. (**F**) Distributions of MurA activity scores from Dewachter et al.(7) for open-biased, closed-biased, and neutral populations. ***p < 0.001; ****p < 0.0001. (**G**) FabZ is an *E. coli* dehydratase involved in peptidoglycan synthesis that switches between a closed conformation and an open, substrate-bound conformation. Structures were generated using AlphaFold2, templating on open and closed structures of *H. pylori* FabZ (PDB: 4ZJB and 2GLL). AS, active site. (**H**) Distributions of FabZ activity scores from Dewachter et al.(7) for closed-biased, open-biased, and neutral populations. *p<0.05; **p<0.01; ****p < 0.0001.

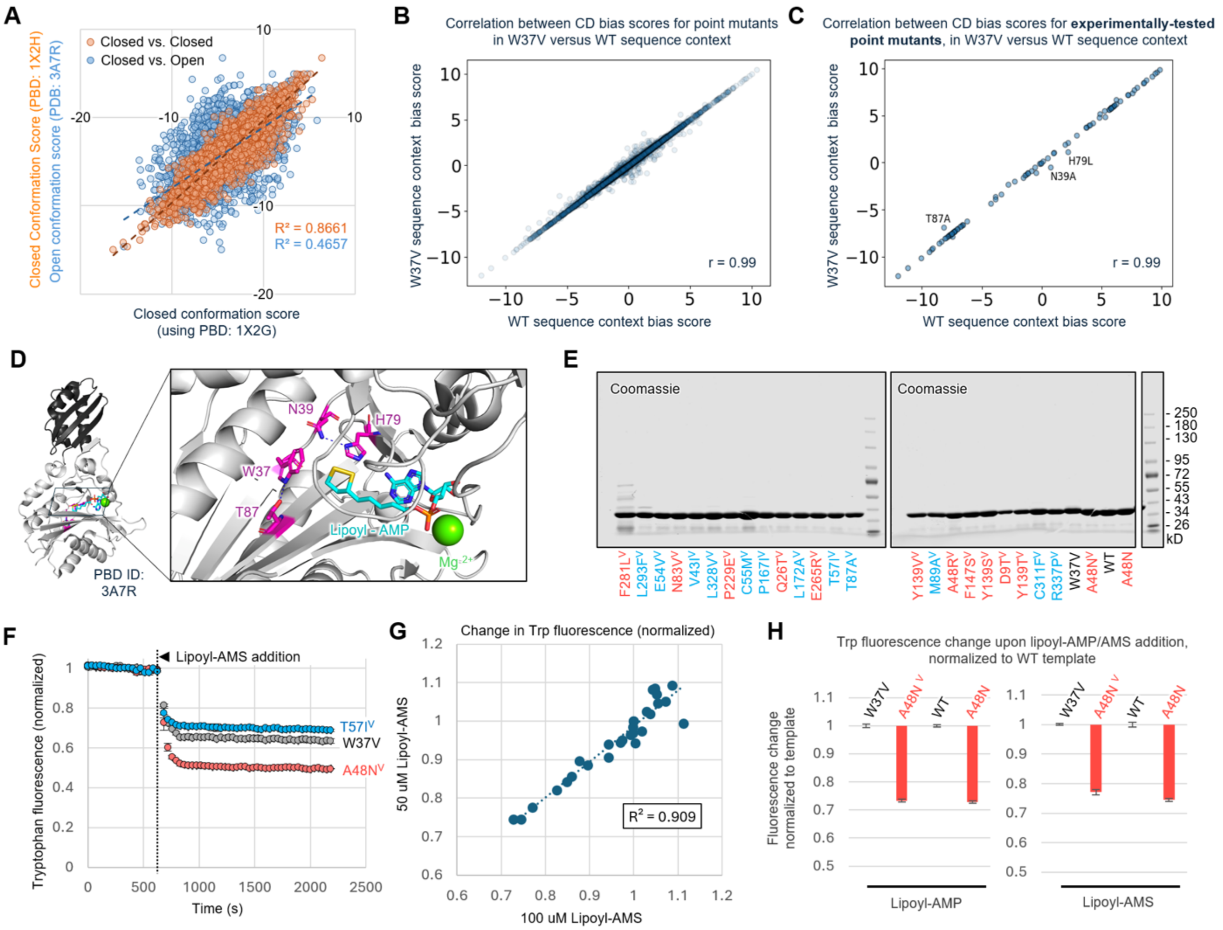

**Figure S4. Additional data related to Figure 3 on LplA.** (**A**) CB analysis using two closed structures of LplA (orange) versus open and closed LplA structures (blue). The former shows a narrower spread and higher correlation. (**B**) Correlation between CB bias scores for point mutants on a wild-type LplA background versus W37V LplA background. Closed and open structures used are PDB 1X2G and 3A7R, respectively. CB bias score is defined as the difference between the ProteinMPNN score on the open structure and the ProteinMPNN score on the closed structure. (**C**) Same as (B) but plotted for the 99 LplA mutants that we experimentally tested in **Figure 4C**. (**D**) Mutations that deviate from the diagonal in (C) are close to the W37 sidechain in the LplA structure. (**E**) Coomassie-stained SDS-PAGE analysis of purified LplA variants used for tryptophan fluorescence measurements in **Figure 3F**. (**F**) Decrease in intrinsic Trp fluorescence over time for W37V-LplA, an open-biased variant, and a closed-biased variant. 100 uM lipoyl-AMS was added at the indicated time. (**G**) Correlation between experiments measuring fold-change in Trp fluorescence for LplA variants, using 50 uM (y axis) or 100 uM (x axis) lipoyl-AMS. (**H**) Trp fluorescence change upon lipoyl-AMP or lipoyl-AMS addition for WT, A48N, W37V, and A48N$^V$ LplA variants.

**A**

| + Lipoyl-AMS Samples | Sample | Closed conf. | Open conformation | | |
|---|---|---|---|---|---|
| | | | Monomer | Dimer | Total Open |
| Closed-biased | C55M$^{V}$ + LA-AMS | 88.2 ± 0.1 % | 11.9 ± 0.1 % | 0 ± 0 % | 11.9 ± 0.1 % |
| | T57I$^{V}$ + LA-AMS | 84 ± 9.1 % | 16.1 ± 9.1 % | 0 ± 0 % | 16.1 ± 9.1 % |
| Template (neutral) | W37V + LA-AMS | 77.6 ± 1.6 % | 22.4 ± 1.6 % | 0 ± 0 % | 22.4 ± 1.6 % |
| | WT + LA-AMS | 77.8 ± 0.2 % | 22.3 ± 0.2 % | 0 ± 0 % | 22.3 ± 0.2 % |
| Open-biased | E265R$^{V}$ + LA-AMS | 10.4 ± 2.6 % | 89.2 ± 2.9 % | 0.6 ± 0.2 % | 89.7 ± 2.7 % |
| | A48N$^{V}$ + LA-AMS | 1 ± 1.7 % | 59.4 ± 10.6 % | 39.6 ± 12.2 % | 99 ± 1.7 % |
| | A48N + LA-AMS | 0 ± 0 % | 58.2 ± 3.4 % | 41.8 ± 3.4 % | 100 ± 0 % |

| Apo Samples | Sample | Closed conf. | Open conformation | | |
|---|---|---|---|---|---|
| | | | Monomer | Dimer | Total Open |
| Closed-biased | C55M$^{V}$ | 95.4 ± 1.5 % | 0 ± 0 % | 4.7 ± 1.5 % | 4.7 ± 1.5 % |
| | T57I$^{V}$ | 95.4 ± 1.3 % | 0 ± 0 % | 4.7 ± 1.3 % | 4.7 ± 1.3 % |
| Template (neutral) | W37V | 96.7 ± 2.6 % | 0 ± 0 % | 3.4 ± 2.6 % | 3.4 ± 2.6 % |
| | WT | 94.3 ± 1.6 % | 0 ± 0 % | 5.7 ± 1.6 % | 5.7 ± 1.6 % |
| Open-biased | E265R$^{V}$ | 97.1 ± 1.1 % | 0 ± 0 % | 2.9 ± 1.1 % | 2.9 ± 1.1 % |
| | A48N$^{V}$ | 98.3 ± 1.1 % | 0 ± 0 % | 1.8 ± 1.1 % | 1.8 ± 1.1 % |
| | A48N | 96.9 ± 1.6 % | 0 ± 0 % | 3.1 ± 1.6 % | 3.1 ± 1.6 % |

**B**

| Sample | Closed conf. | Open conformation | | |
|---|---|---|---|---|
| | | Monomer | Dimer | Total Open |
| Wild-type + lipoic acid | 93% | 0% | 7% | 7% |
| Wild-type + lipoyl-AMP | 75% | 25% | 0% | 25% |
| A48N LplA + lipoic acid | 95% | 0% | 5% | 5% |
| A48N LplA + lipoyl-AMP | 0% | 39% | 61% | 100% |

**C**

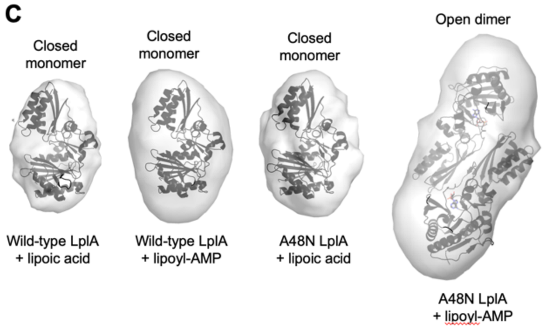

**D**

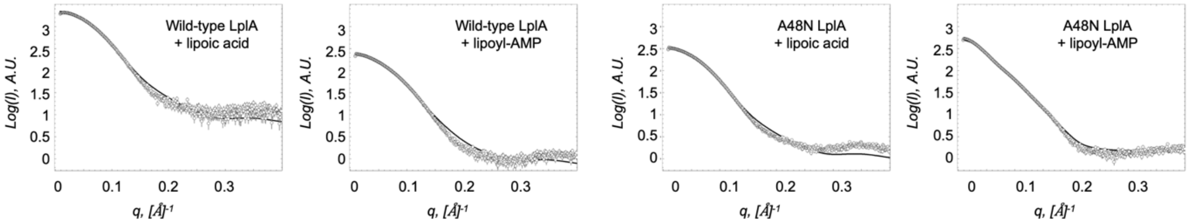

**E**

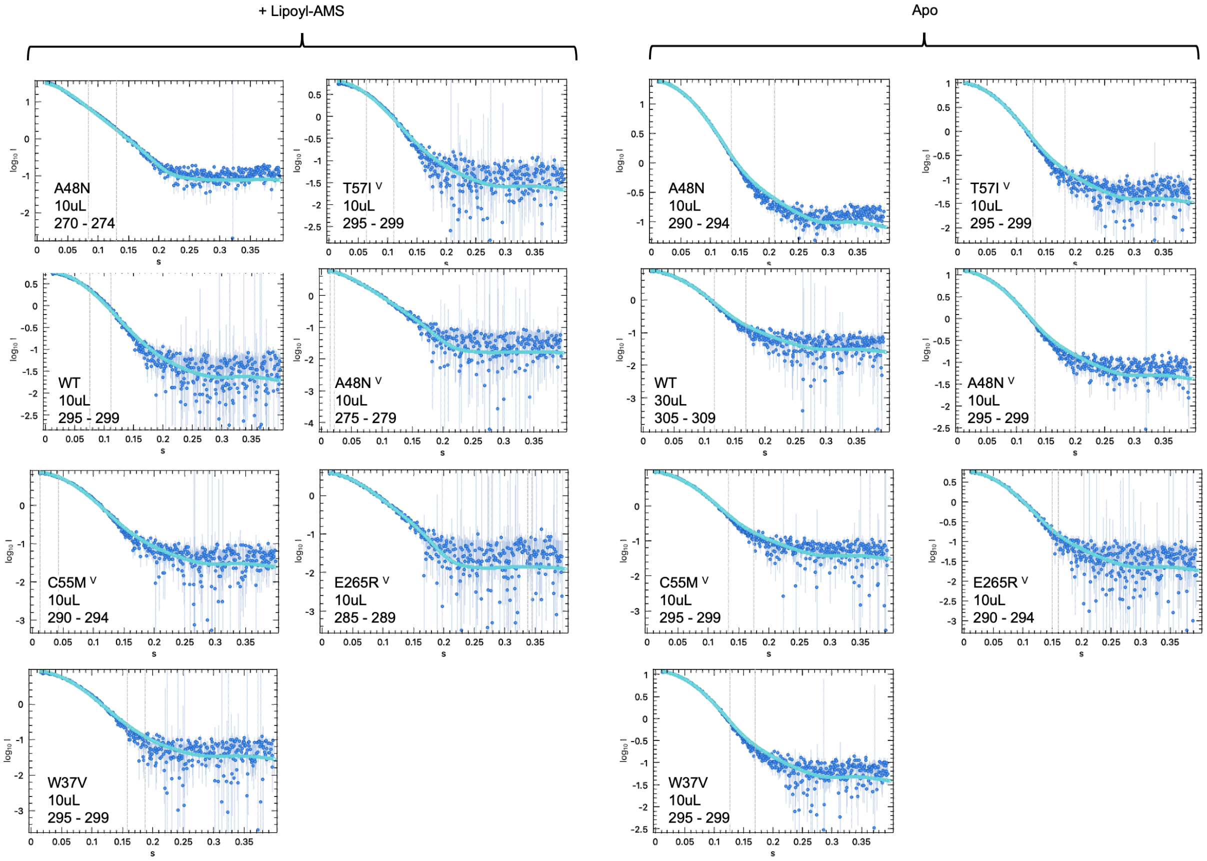

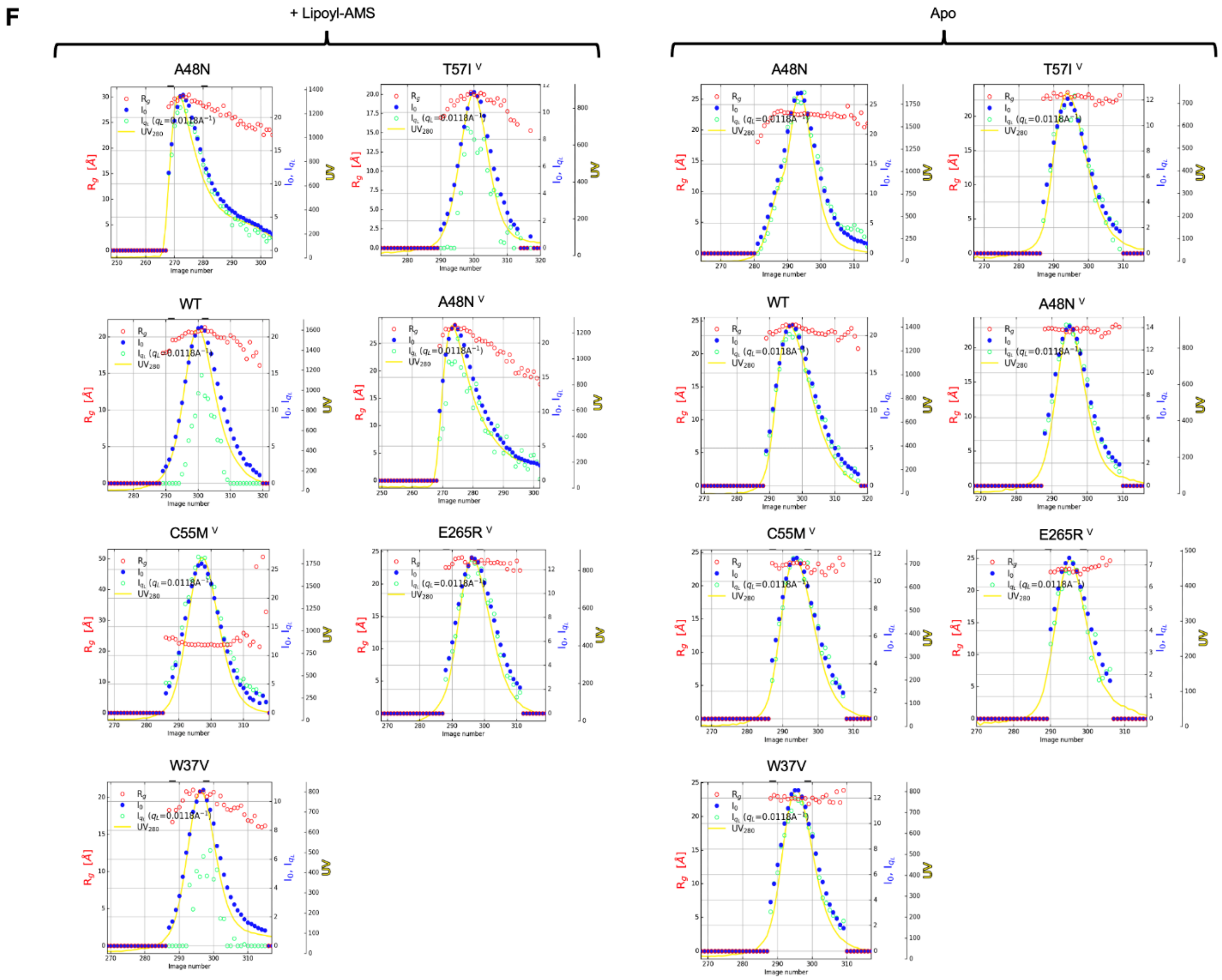

**Figure S5. Data related to SEC-SAXS analysis of LplA variants.** (**A**) Summary of fractional occupancies predicted by Oligomer, for LplA variants alone (apo form, bottom) or after binding to lipoyl-AMS (top). Each condition was repeated 2-5 times. Errors, ±1 std. dev. (**B**) Same as (A), but comparing lipoic acid- and lipoyl-AMP bound conditions. Each condition was repeated 2-5 times. Errors, ±1 std. dev. (**C**) DENSS *ab initio* modeling of protein envelopes predicted for samples in (B), fit with the major LplA conformer detected under each condition. The open dimer structure was generated by AlphaFold3. (**D**) Kratky plots for samples in (B) showing log(**I**) versus q for oligomer fit (black line) to experimental data (grey circles), assuming a mixture of three species: open monomer (3A7R), closed monomer (1X2H), and open dimer (AlphaFold3 prediction). For the WT + lipoic acid sample, evolving factor analysis was used to extract the monomeric fraction (see Methods). (**E**) Representative SAXS scattering curves for samples in (A) and **Figure 3D**, showing log(I) versus q for oligomer fit (light blue line) to experimental data (dark blue circles), assuming a mixture of three species: open monomer (3A7R), closed monomer (1X2H), and open dimer (AlphaFold3 prediction). For select samples, evolving factor analysis was used to extract the monomeric fraction (see Methods). Each plot is labeled with LplA variant, injection volume, and image number window analyzed. (**F**) Representative SEC elution profiles for samples in (A) and **Figure 3D**. X axis is image number, and y axis shows radius of gyration (red circles), UV absorbance (yellow line), $I_{QL}$ (green circles), and $I_o$ (blue circles). Data shown are representative of 2-4 replicates per condition.

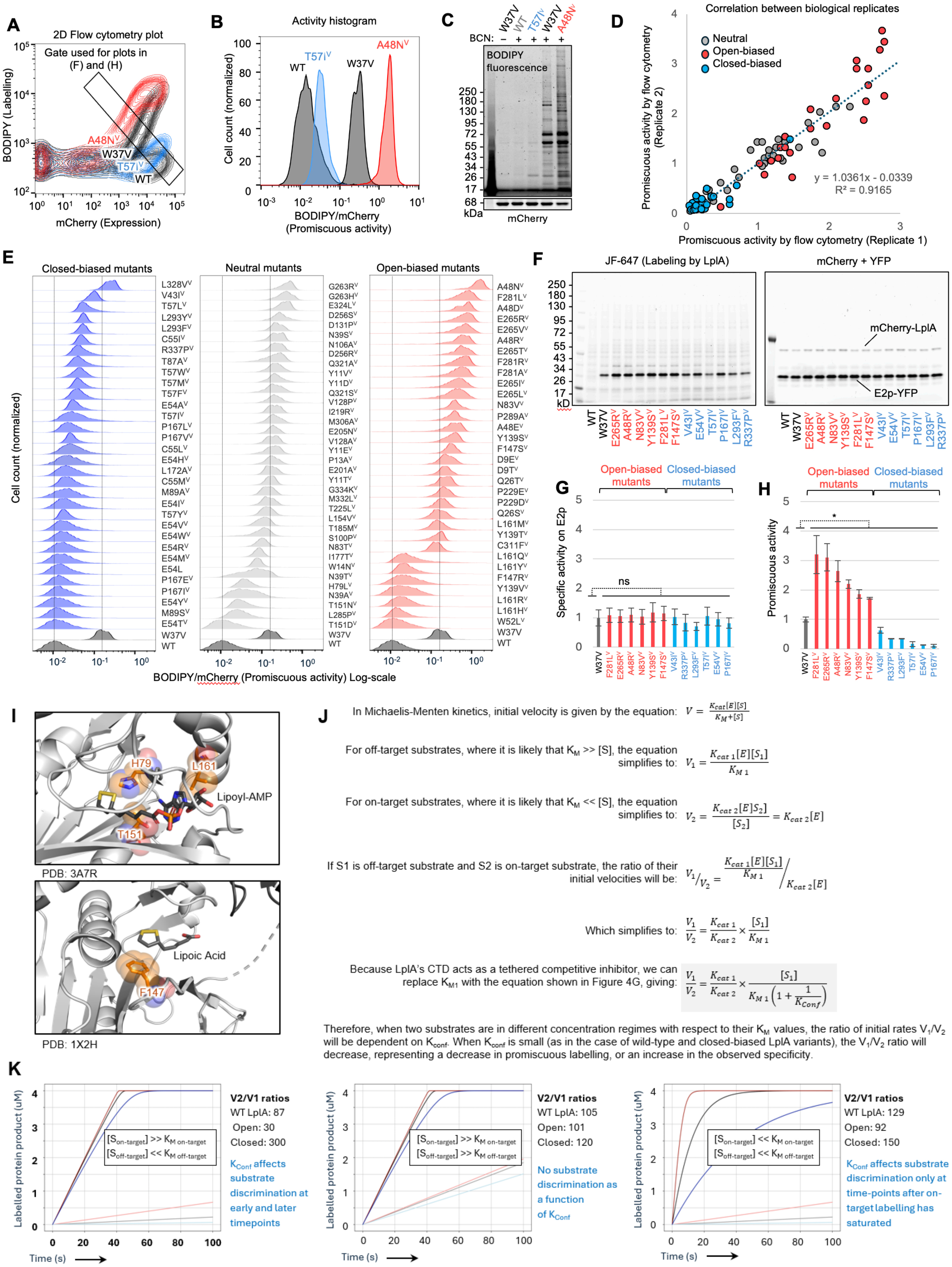
A
2D Flow cytometry plot
Gate used for plots in (F) and (H)
BODIPY (Labelling)
mCherry (Expression)
A48N^V
W37V
T57I^V
WT
B
Activity histogram
Cell count (normalized)
WT
T57I^V
W37V
A48N^V
BODIPY/mCherry (Promiscuous activity)
C
BCN:
W37V
WT
T57I^V
W37V
A48N^V
BODIPY fluorescence
mCherry
kDa
D
Correlation between biological replicates
Neutral
Open-biased
Closed-biased
Promiscuous activity by flow cytometry (Replicate 2)
Promiscuous activity by flow cytometry (Replicate 1)
y = 1.0361x - 0.0339
R² = 0.9165
E
Closed-biased mutants
Neutral mutants
Open-biased mutants
Cell count (normalized)
BODIPY/mCherry (Promiscuous activity) Log-scale
F
JF-647 (Labeling by LplA)
mCherry + YFP
mCherry-LplA
E2p-YFP
kD
G
Open-biased mutants
Closed-biased mutants
Specific activity on E2p
ns
H
Open-biased mutants
Closed-biased mutants
Promiscuous activity
I
H79
L161
Lipoyl-AMP
T151
PDB: 3A7R
Lipoic Acid
F147
PDB: 1X2H
J
In Michaelis-Menten kinetics, initial velocity is given by the equation: $V = \frac{K_{cat}[E][S]}{K_M+[S]}$
For off-target substrates, where it is likely that $K_M >> [S]$, the equation simplifies to: $V_1 = \frac{K_{cat\,1}[E][S_1]}{K_{M\,1}}$
For on-target substrates, where it is likely that $K_M << [S]$, the equation simplifies to: $V_2 = \frac{K_{cat\,2}[E][S_2]}{[S_2]} = K_{cat\,2}[E]$
If S1 is off-target substrate and S2 is on-target substrate, the ratio of their initial velocities will be: $V_1/V_2 = \frac{K_{cat\,1}[E][S_1]}{K_{M\,1}} \Big/ K_{cat\,2}[E]$
Which simplifies to: $\frac{V_1}{V_2} = \frac{K_{cat\,1}}{K_{cat\,2}} \times \frac{[S_1]}{K_{M\,1}}$
Because LplA's CTD acts as a tethered competitive inhibitor, we can replace $K_{M1}$ with the equation shown in Figure 4G, giving: $\frac{V_1}{V_2} = \frac{K_{cat\,1}}{K_{cat\,2}} \times \frac{[S_1]}{K_{M\,1}\left(1+\frac{1}{K_{Conf}}\right)}$
Therefore, when two substrates are in different concentration regimes with respect to their $K_M$ values, the ratio of initial rates $V_1/V_2$ will be dependent on $K_{conf}$. When $K_{conf}$ is small (as in the case of wild-type and closed-biased LplA variants), the $V_1/V_2$ ratio will decrease, representing a decrease in promiscuous labelling, or an increase in the observed specificity.
K
Labelled protein product (uM)
Time (s)
[S_on-target] >> K_M on-target
[S_off-target] << K_M off-target
V2/V1 ratios
WT LplA: 87
Open: 30
Closed: 300
K_Conf affects substrate discrimination at early and later timepoints
[S_on-target] >> K_M on-target
[S_off-target] >> K_M off-target
V2/V1 ratios
WT LplA: 105
Open: 101
Closed: 120
No substrate discrimination as a function of K_Conf
[S_on-target] << K_M on-target
[S_off-target] << K_M off-target
V2/V1 ratios
WT LplA: 129
Open: 92
Closed: 150
K_Conf affects substrate discrimination only at time-points after on-target labelling has saturated

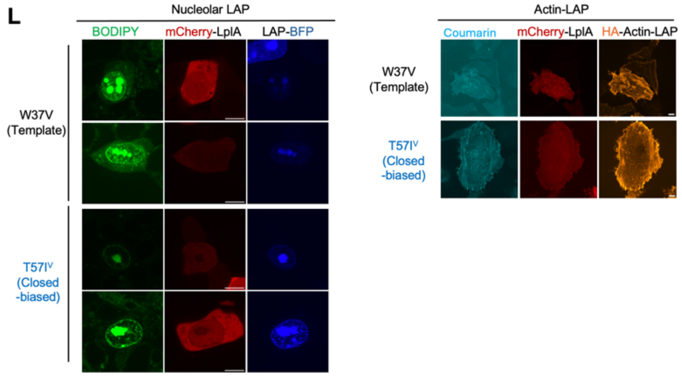

**Figure S6. Additional data related to LplA Figure 4.** (**A**) Representative 2D flow cytometry plots for assay shown in **Figure 4A**. A48N$^{V}$ is open-biased and T57I$^{V}$ is closed-biased. Wild-type (WT) LplA is unable to use BCN as a substrate. (**B**) Conversion of 2D flow cytometry data into 1D promiscuous activity histograms, using the rectangular gate in (A). (**C**) SDS-PAGE analysis of HEK lysates from (A) showing promiscuous labeling of many endogenous proteins for the open-biased variant A48N$^{V}$ and a moderate degree of promiscuity for W37V-LplA. (**D**) Correlation between biological replicates for experiment in **Figure 4C**. (**E**) Data from **Figure 4C** plotted on a log rather than linear scale. (**F**) SDS-PAGE and in-gel fluorescence imaging of LplA's sequence-specific labeling activity. HEK cells expressing LplA variant and the LplA acceptor protein E2p-YFP in a 1:50 ratio were labeled with BCN for 5 minutes before lysis and Click with JF646-tetrazine. This experiment was performed three times. (**G**) Quantitation of data in (F). ns, not significant. (**H**) Promiscuous activity of variants as measured by flow cytometry data in **Figure 4C**, normalized to the template W37V-LplA. Three replicates. Errors, ±1 std. dev. * $p<0.05$. (G) and (H) were used to generate the bar graph in **Figure 4F**. (**I**) LplA sidechains predicted to clash with the BCN substrate when mutated by CB. These variants are marked with X in **Figure 5C, H, S9A-D**. (**J**) Kinetic model of LplA. Under the assumption that on-target substrates have low $K_M$ (<< [$S_{on\text{-}target}$]) and off-target substrates have high $K_M$ (>> [$S_{off\text{-}target}$]), we find that the ratio of their initial velocities is dependent on $K_{Conf}$. (**K**) Modelling of LplA kinetics under varying concentration to $K_M$ ratios for cognate and noncognate substrates. Only when one substrate is in excess of its $K_M$ and another substrate is below its $K_M$ does the ability of LplA to discriminate between these substrates change with $K_{Conf}$. Exact modeling conditions outlined in **Methods**. (**L**) Additional fields of view for site-specific LAP fusion protein labeling in **Figure 4I**. Scale bars, 10 um.

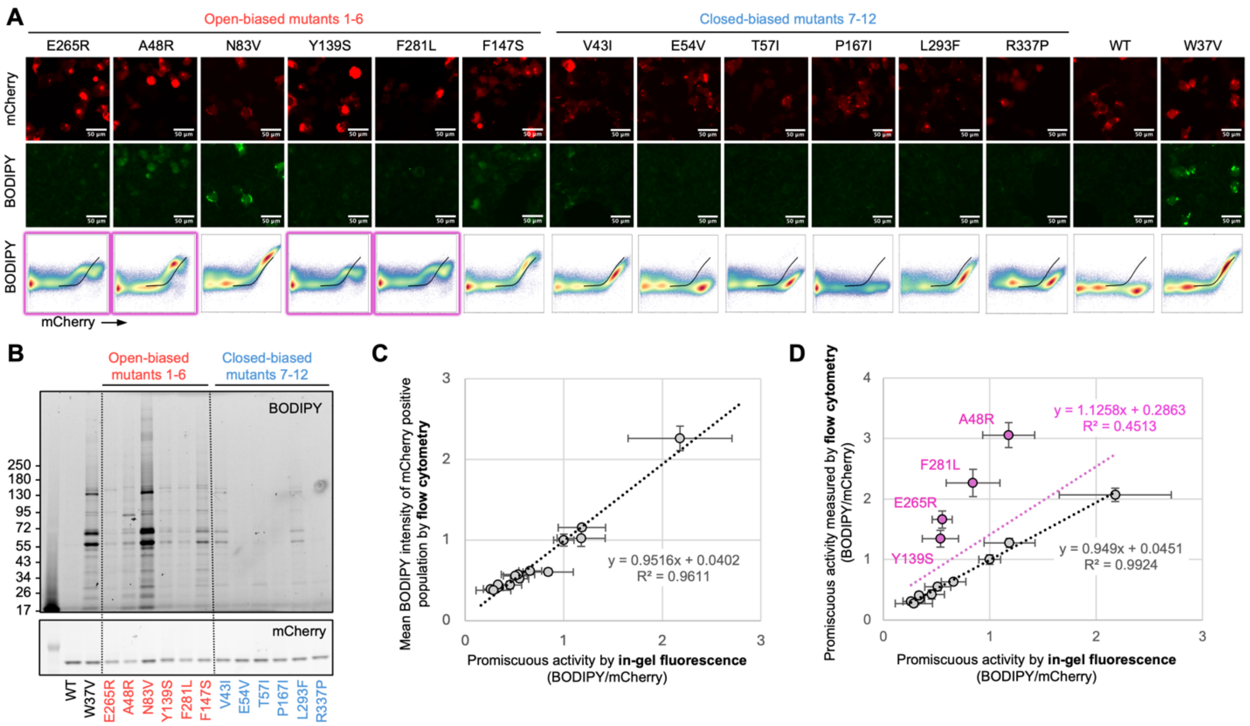

**Figure S7. Correlation between three different assays of LplA promiscuous activity: flow cytometry, fluorescence microscopy, and SDS-PAGE.** See **Supporting Text** for discussion of these assays. All assays were performed on the template W37V LplA and 6 open-biased and 6-closed biased variants characterized by Trp fluorescence in **Figure 3F.** (**A**) Confocal imaging of fixed HEK 293T cells after labeling with BCN and tetrazine-BODIPY as in **Figure 4A**. Flow cytometry plots for the same samples are shown below. Mutants that show BODIPY saturation at high mCherry-LplA expression levels are outlined in pink. (**B**) SDS-PAGE and in-gel fluorescence imaging of lysates from samples in (A). (**C**) Correlation between flow cytometry data from (A) and in-gel fluorescence data from (B). On the y-axis, mean BODIPY signal from all mCherry-positive cells was quantified, and normalized to that of W37V. (**D**) Same correlation analysis as in (C) but using BODIPY/mCherry ratio from the rectangular gate in **Figure S6A** (values are normalized to that of W37V). Using this method of quantifying flow cytometry data, we observe poor correlation with SDS-PAGE for open-biased variants that show BODIPY saturation (pink mutants in (A)).

Open-biased LplA variants

**A**

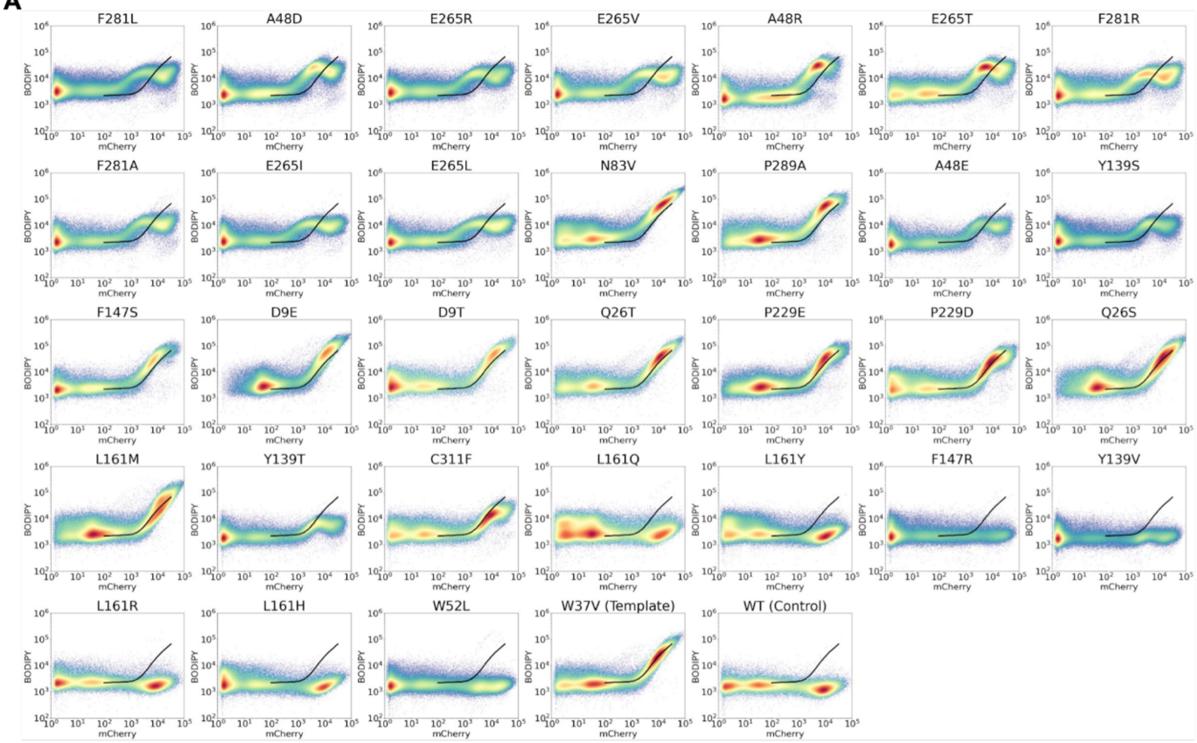

**B**

Neutral LplA variants

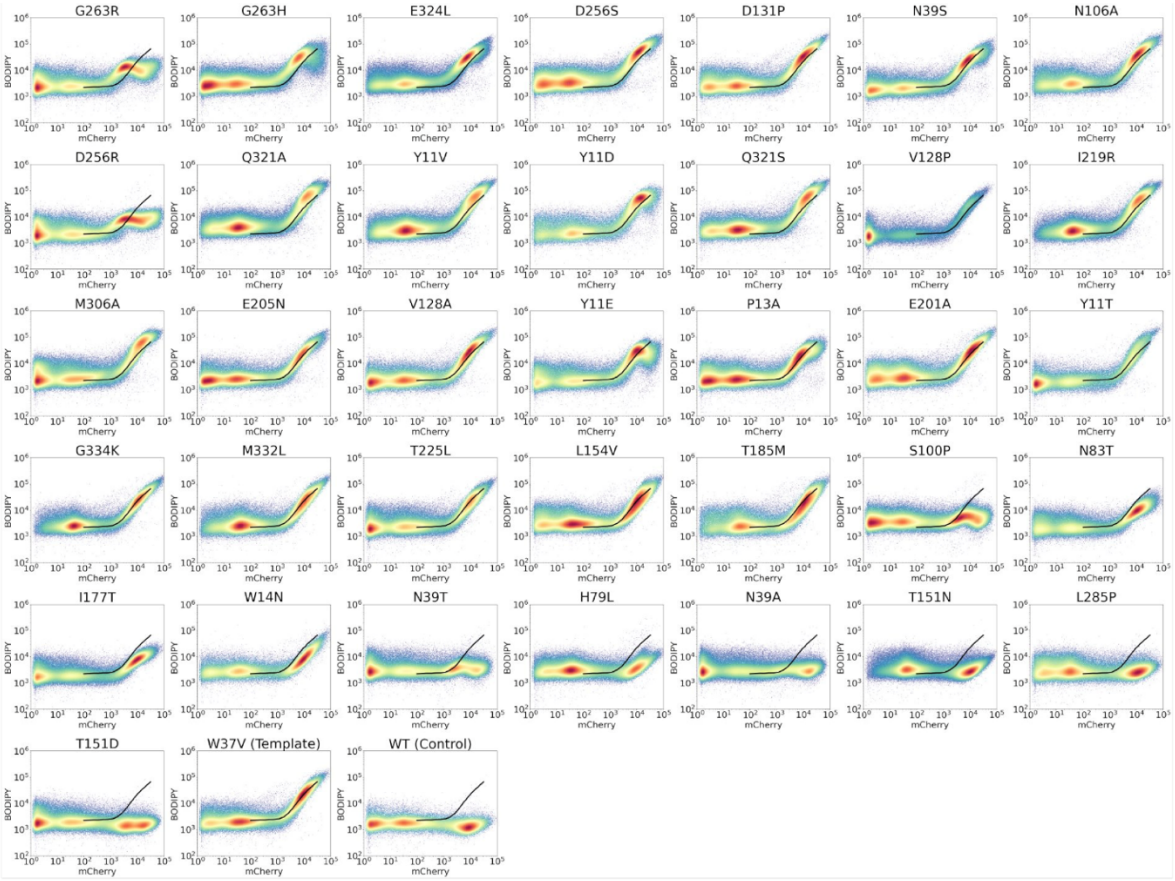

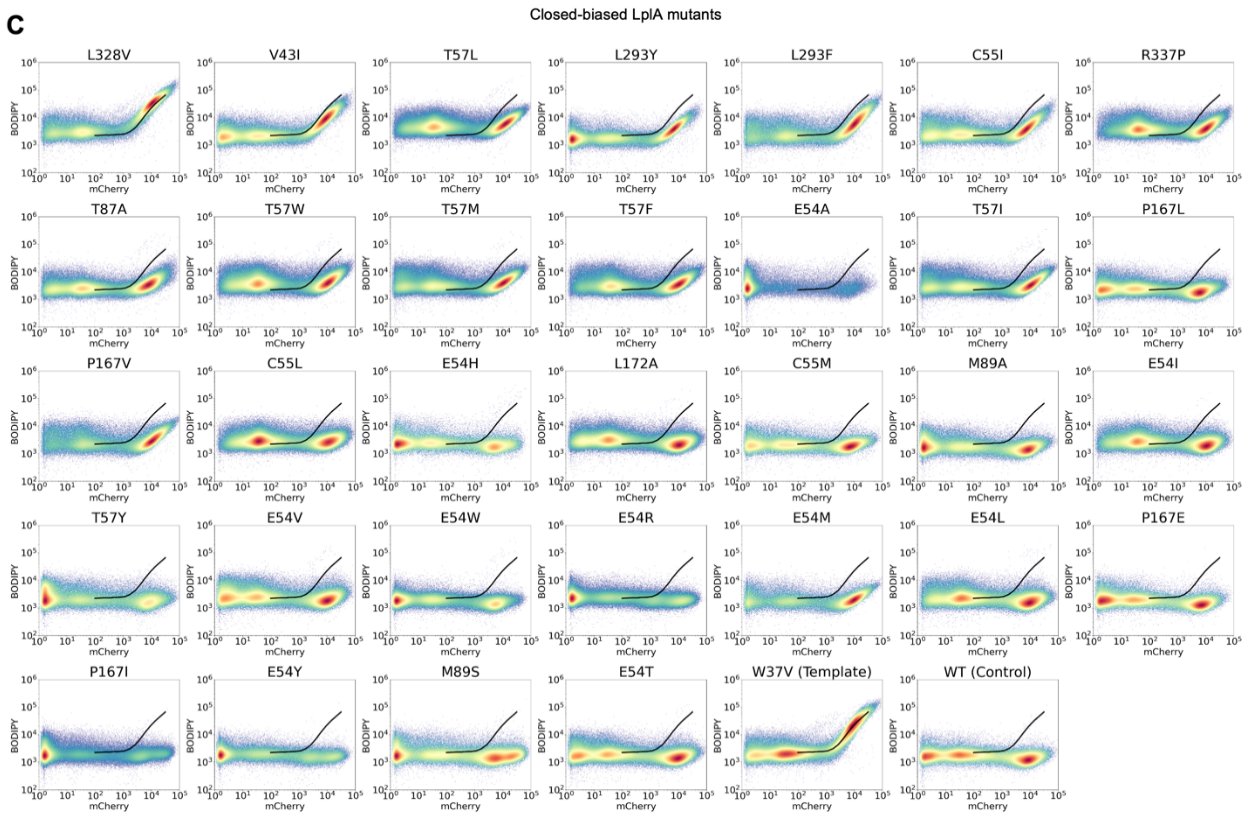

**Figure S8. 2D Flow cytometry plots for all LplA variants.** All variants are on a W37V background except for WT. (**A**) Open-biased variants (**B**) neutral variants, and (**C**) closed-biased variants. The black reference line in all plots shows the mean BODIPY vs. mCherry signal for the W37V-LplA template.

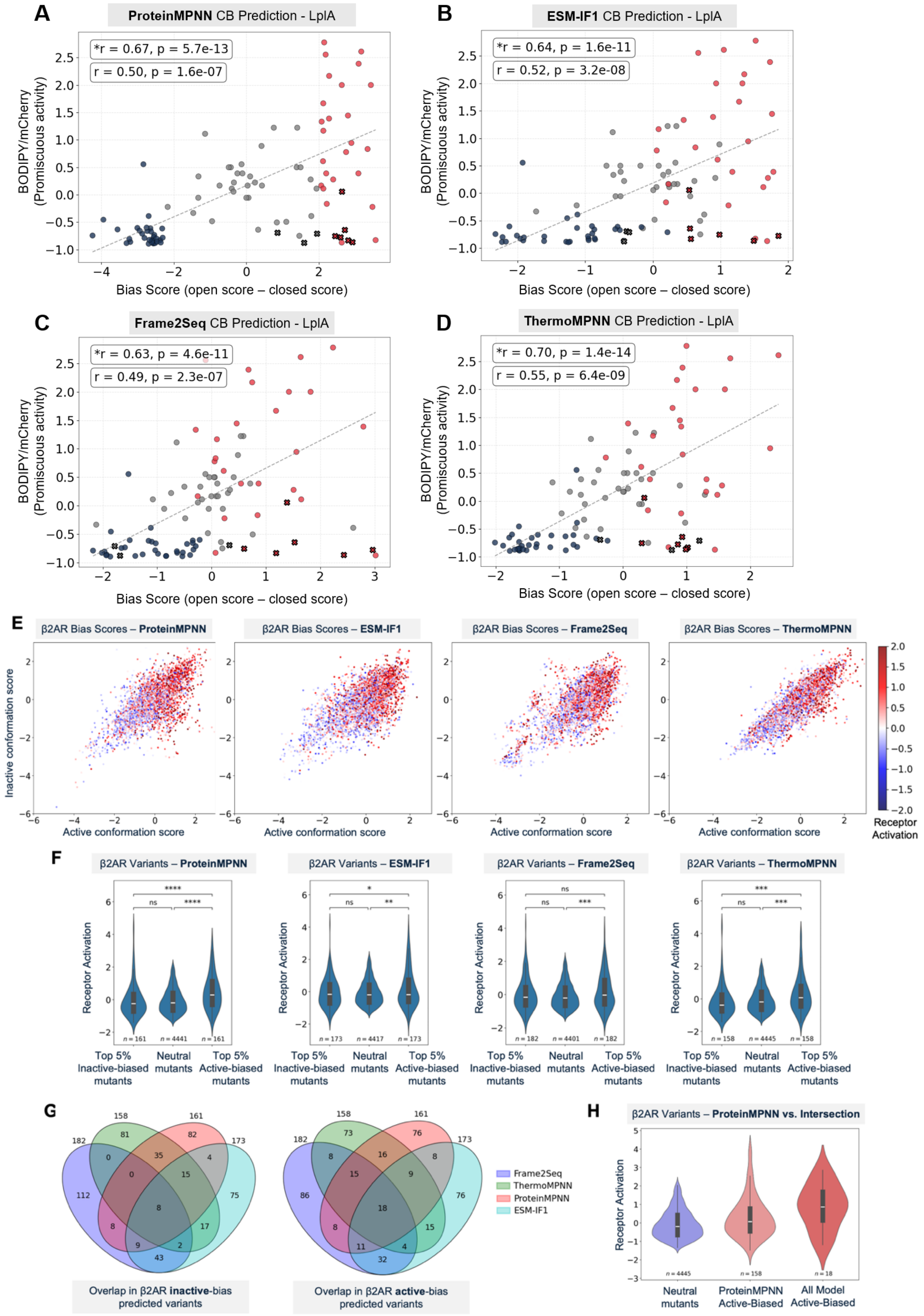

A
ProteinMPNN CB Prediction - LplA
*r = 0.67, p = 5.7e-13
r = 0.50, p = 1.6e-07
BODIPY/mCherry (Promiscuous activity)
Bias Score (open score – closed score)
B
ESM-IF1 CB Prediction - LplA
*r = 0.64, p = 1.6e-11
r = 0.52, p = 3.2e-08
BODIPY/mCherry (Promiscuous activity)
Bias Score (open score – closed score)
C
Frame2Seq CB Prediction - LplA
*r = 0.63, p = 4.6e-11
r = 0.49, p = 2.3e-07
BODIPY/mCherry (Promiscuous activity)
Bias Score (open score – closed score)
D
ThermoMPNN CB Prediction - LplA
*r = 0.70, p = 1.4e-14
r = 0.55, p = 6.4e-09
BODIPY/mCherry (Promiscuous activity)
Bias Score (open score – closed score)
E
β2AR Bias Scores – ProteinMPNN
β2AR Bias Scores – ESM-IF1
β2AR Bias Scores – Frame2Seq
β2AR Bias Scores – ThermoMPNN
Inactive conformation score
Active conformation score
Receptor Activation
F
β2AR Variants – ProteinMPNN
β2AR Variants – ESM-IF1
β2AR Variants – Frame2Seq
β2AR Variants – ThermoMPNN
Receptor Activation
Top 5% Inactive-biased mutants
Neutral mutants
Top 5% Active-biased mutants
G
Frame2Seq
ThermoMPNN
ProteinMPNN
ESM-IF1
Overlap in β2AR inactive-bias predicted variants
Overlap in β2AR active-bias predicted variants
H
β2AR Variants – ProteinMPNN vs. Intersection
Receptor Activation
Neutral mutants
ProteinMPNN Active-Biased
All Model Active-Biased

**Figure S9. Testing different inverse folding models for CB scoring.** (**A**)-(**D**) Correlation between LplA promiscuous labeling activities and CB bias scores calculated using ProteinMPNN, ESM-IF1, Frame2Seq, or ThermoMPNN. (**E**) CB plots for β2AR variants, scored using the indicated IFM. Points are colored by experimentally-determined receptor variant activity(4). (**F**) Comparison of top 5% active/inactive-biased β2AR variants determined by CB scoring with (from left to right) ProteinMPNN, ESM-IF1, Frame2Seq, or ThermoMPNN. *p<0.05, **p<0.01, ***p<0.001, ****p<0.0001 (**G**) 4-way Venn diagram of overlapping active-biased and inactive-biased predictions (top 5%) by each model. (**H**) Distribution of receptor activities for conformationally neutral mutants, ProteinMPNN-predicted active-biased variants, and variants predicted as active-biased by all four models.

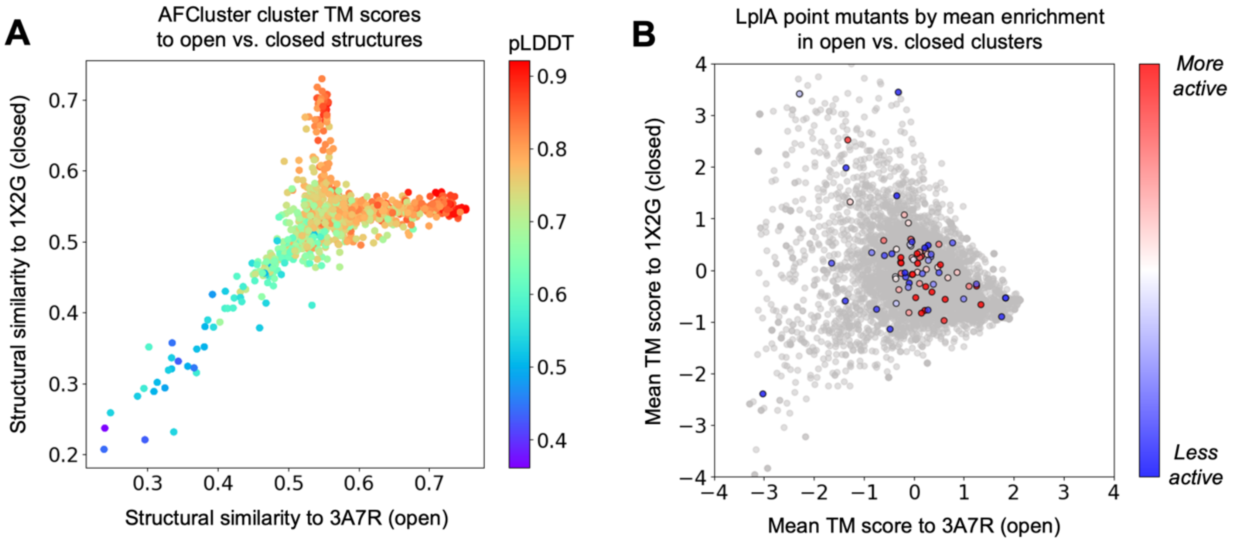

**C**

LplA mutants unable to be scored by AFCluster

| Open mutations (n = 31) | Neutral mutations (n=36) | Closed mutations (n = 32) | All mutations (n = 6403) |
|---|---|---|---|
| A48D, A48R, A48E, L161Q, L161Y, L161H | S100P, N39A, T151N | L293Y, R337P, E54A, E54H, E54I, E54W, E54R, E54M, E54L, E54Y, E54T | 1250 point mutations not present in MSA |

**Figure S10. Benchmarking CB against AFCluster.** (**A**) Plot showing structural similarity of clusters generated by AFCluster to LplA open (PDB: 3A7R) and closed (PBD: 1X2G) conformations. Each point represents a cluster of evolutionarily-related sequences from the LplA multiple sequence alignment (MSA). Higher template modeling score (TM-score) means higher structural similarity to the indicated structure. (**B**) Plot of LplA point mutations present in the MSA, showing mean structural similarity to open (3A7R) and closed (1X2G) conformations. Mutations absent in the MSA (see (C)) were not plotted. Experimentally tested LplA mutations are colored by their promiscuous activity scores from **Figure 4C**. (**C**) Table summarizing LplA point mutations not present in the MSA. These variants can be scored using CB but not using AFCluster.

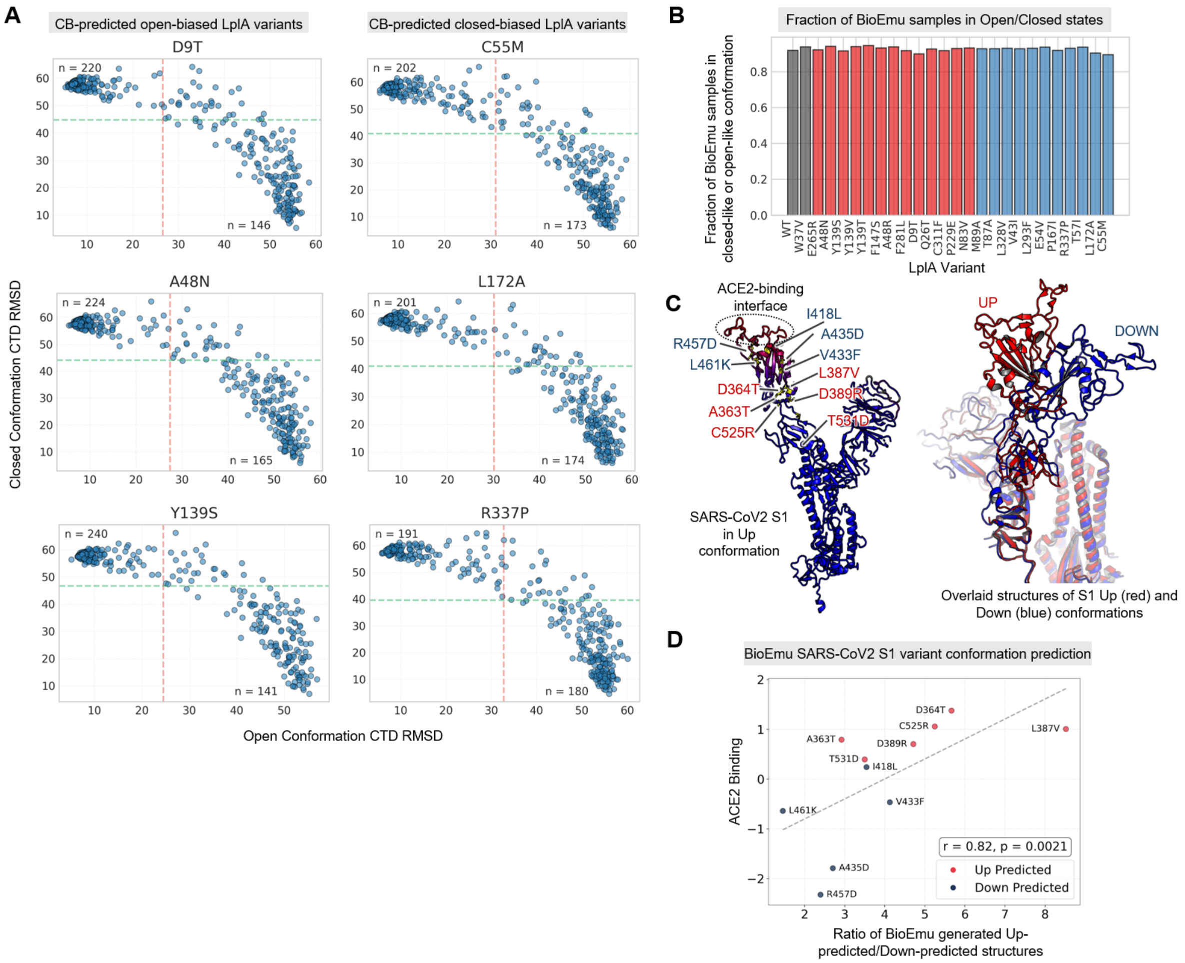

**Figure S11. Benchmarking CB against BioEmu.** (**A**) Detail for sampled structures generated by BioEmu for six CB-predicted open-biased (left) and closed-biased (right) variants. The position of each sampled structure in the graph represents aligned CTD (C-terminal domain) distance to open vs. closed structures. (**B**) Percentage of structures sampled by BioEmu that are in an open-like or closed-like states for LplA. (**C**) CB-predicted SARS-CoV2 S1 down and up-biased variants selected for analysis by BioEmu(8). Red-colored mutants show increased ACE2 binding in Starr et al.(2), and blue mutants show decreased ACE2 binding. Red and blue mutants were all predicted by CB. Right: Overlaid structures of S1 in up (red) and down (blue) conformations. (**D**) BioEmu was used to generate a conformational ensemble for each variant from (C). Sampled conformations were assigned to up/down conformations, and ratio of “up” occupancy vs. “down” occupancy is shown for each variant plotted against ACE2 binding.

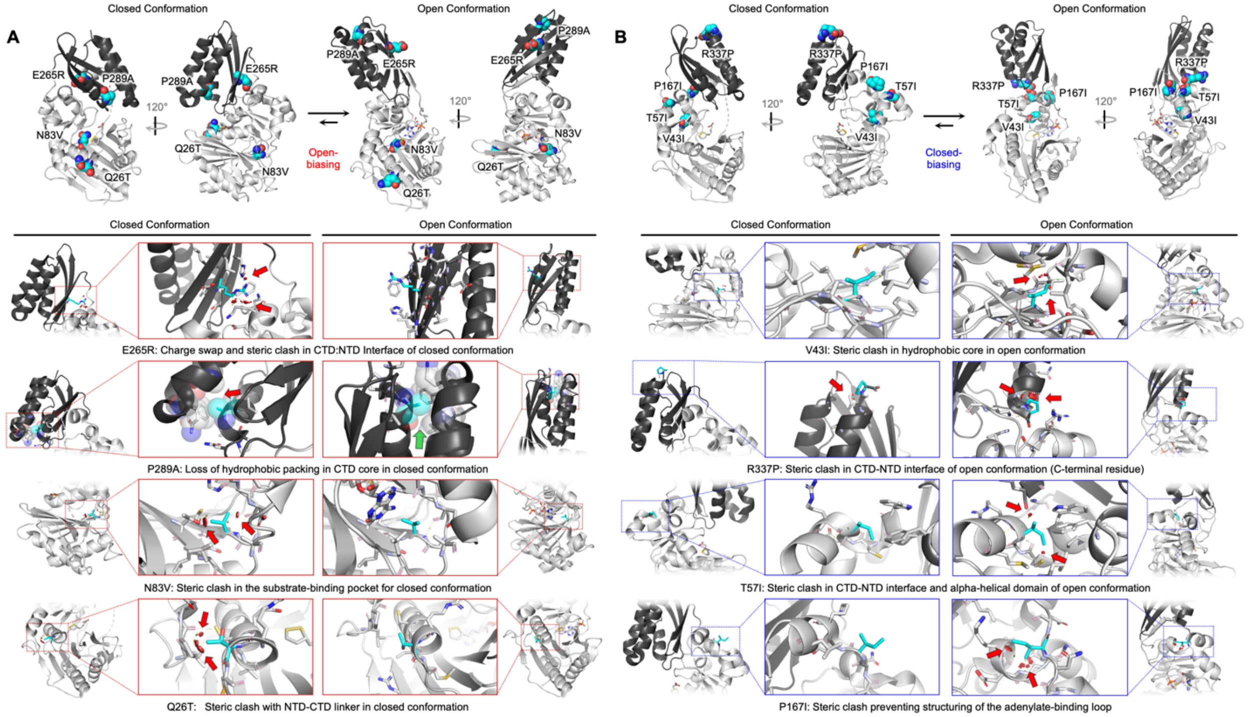

**Figure S12. Diversity of conformation-biasing mutations in LplA.** Structural analysis of four open-biased (**A**) and four closed-biased (**B**) LplA variants. Mutations are modeled into the closed (1X2G) and open (3A7R) structures of LplA using PyMOL. Mutations are observed at the interface between NTD and CTD (e.g. A48, E265, F281, R337, T57I), in the hydrophobic core (P289, L293, V43I), in the substrate-binding loops / beta-sheet (F147, N83, Y139, P167), and on the exterior surface of the protein (D9, Q26). Several open-biased variants show clashes in the closed conformation, which are alleviated in the open conformation, and vice versa for closed-biased variants. For P289A (left, second row), a loss of hydrophobic packing in the CTD is predicted for the closed conformation but not open conformation.

**Supplementary Table 1.** CB scores for LplA, generated using ProteinMPNN, ESM-IF1, Frame2Seq, and ThermoMPNN on the open and closed structures of LplA (3A7R and 1X2G, respectively). Promiscuous activity (median BODIPY/mCherry for the gated population of cells) is also given for the LplA variants that were experimentally evaluated.

**Supplementary Table 2.** Tryptophan fluorescence data for LplA, used in **Figure 3F**.

**Supplementary Table 3-9.** DMS datasets with CB scoring for K-Ras, SARS-CoV2, β2AR, Src, B-Raf, MurA, and FabZ.

### Supporting Text

**Considering just the subset of variants with above-average scores on both backbone conformations ("Quadrant 1" variants, Figure S1).** We used the K-Ras dataset to ask whether CB's effects require destabilization of one conformation or if effects can also be seen from neutral or stabilizing mutations only. To answer this, we repeated the K-Ras analysis using just a stabilizing subset of variants with above-average scores on both State 1 and State 2 conformations (quadrant 1, or Q1 variants in **Figure 2B**). These variants show above-average expression compared to wild-type K-Ras(1), suggesting that they are as stable or more stable than the template (**Figure S1B**). **Figure S1C** shows that even when analyzing the Q1 subset, the same highly significant effect is seen across all effector datasets for K-Ras, albeit with slightly reduced effect size and significance. When we performed the same analysis on K-Ras single mutants only (**Figures S1E-F**), the significance of the effect was again reduced for Q1 variants, but a binding effect was still seen for 4/5 effector datasets for which an effect was observed across all single mutants. Thus, we conclude that variants that destabilize one conformation while stabilizing the other (Q2 and Q4 variants in **Figure 2B**) are likely to produce the strongest effect on conformational occupancy, but especially in the context of multiple mutations, a strong effect can be seen even when selecting for only stabilizing mutations. Importantly, we observed that Q2/Q4 variants have on average slightly above-wild-type expression (**Figure S1B**), and thus do not necessarily compromise the overall stability of the protein, which would reduce the utility of CB for engineering purposes. We also performed a similar Q1 analysis of the SARS-CoV2 spike and β2AR DMS datasets and reached similar conclusions (**Figure S1G-J**).

**Flow cytometry assay for quantification of promiscuous labeling activity (Figure S7).** To quantify the promiscuous labeling activity of W37V LplA and its CB-designed point mutants, we opted to use a flow cytometry assay rather than the more traditional western blot readout. This assay is faster, and for reasons explained below, also more accurate for W37V LplA-based mutants. We have observed that the W37V mutant is destabilized relative to wild-type LplA, and many point mutants of W37V LplA are even more destabilized, showing a tendency to aggregate at high expression levels inside cells. This is apparent from imaging of mCherry-LplA fusions in **Figure S7A**, in which a number of variants show large intracellular aggregates, especially at high LplA expression level. Because these aggregates exhibit low BODIPY labeling, they are likely to be enzymatically inactive. Western blot and in-gel fluorescence analysis is performed on 0.5-1 million cells and averages across all their behaviors. Cells with higher mCherry-LplA expression and inactive aggregates are mixed together with cells that have lower mCherry-LplA expression, even though they exhibit different labeling activities. Averaging across these populations leads to overestimation of active enzyme expression and underestimation of activity for unstable variants.

For this reason, we favored flow cytometry as a readout of promiscuous labeling activity (via BODIPY labeling) because it is a *single cell* measurement, and we can gate specifically on lower mCherry-LplA expression levels where protein aggregation is minimal. As shown in **Figure S6A**, our rectangular gate excludes high mCherry-expressing cells with significant aggregation and consistently captures the linear region of the 2D flow plots, where BODIPY signal increases proportionally to mCherry expression. Imaging shows that in this regime, LplA is mostly soluble and active (**Figure S7A**). In **Figure S6B**, we convert 2D flow data into 1D BODIPY/mCherry histograms, reflecting the expression-normalized activity of each enzyme variant. $T57I^V$ and $A48N^V$ are representative closed-biased and open-biased mutants of LplA.

2D flow cytometry plots for all LplA variants are shown in **Figure S8**. The W37V template, and some point mutants, such as N83V, show an extended linear relationship between mCherry (LplA expression level) and BODIPY signal (promiscuous labeling), suggestive of minimal aggregation.

However, many mutants, especially the open-biased ones, show saturation behavior, where BODIPY labeling plateaus at high mCherry expression levels (**Figure S7A**, pink highlighted variants), consistent with aggregation of mCherry-LplA into an inactive fraction. Indeed, curve shape is indicative of LplA stability and aggregation tendency, and mutants with saturated flow curves tend to show more mCherry aggregates by microscopy (**Figure S7A**). It is interesting to note that our "neutral mutants", which have high MPNN scores for both open and closed LplA conformations, tend to show the most linear (non-saturated) flow plots (**Figure S8**), as well as the most smooth, non-punctate expression patterns by mCherry imaging, a testament to ProteinMPNN's ability to predict/design protein stability.

To gain confidence in flow cytometry as a readout of promiscuous activity, for the 6 open-biased and 6 closed-biased mutants analyzed by flow cytometry in **Figure S7A**, we lysed the cells after labeling and analyzed the lysates by SDS-PAGE and in-gel fluorescence (**Figure S7B**). As expected, promiscuous BODIPY tagging of many endogenous proteins of various sizes is observed, and is higher for open-biased variants than for closed-biased ones. Furthermore, **Figure S7C** shows that total BODIPY signal in each gel lane is well-correlated to mean BODIPY signal (for mCherry+ cells) in the 2D flow plots. Thus, it is highly likely that almost all BODIPY signal detected by flow cytometry comes from promiscuous labeling of endogenous proteins (rather than, for example, nonspecific fluorophore accumulation inside cells). Together, these observations support the use of flow cytometry to quantify the relative promiscuous labelling activities of LplA variants.

### Methods

**Conformational Biasing (CB) workflow for deep mutational scanning (DMS) datasets.** We define the score of a protein sequence on a given structure as the following:

$$\text{Score}_{CB}(\text{mut}, \text{wt}) = \sum_{i=1}^{n} \log\left(\frac{P(AA_{i,\text{mut}} \mid \text{Backbone}, AA_{\sim i,\text{mut}})}{P(AA_{i,\text{wt}} \mid \text{Backbone}, AA_{\sim i,\text{wt}})}\right)$$

The score is a pseudo-log-likelihood conditioned on the structure backbone coordinates. In the above equation, $n$ represents the length of the protein sequence, *mut* represents the mutant sequence, *wt* represents the wild-type sequence, and $AA_{\sim i}$ represents all other amino acids other than the one at the *i*-th position. In the standard CB workflow, all point mutants to the starting sequence are scored using ProteinMPNN. The ColabDesign implementation of the ProteinMPNN model in JAX was used. Resulting scores are scaled to a mean of 0 and standard deviation of 1 per structure. For a protein with two states of interest, State *A* and State *B*, State A bias score was calculated by taking the scaled score of State A and subtracting the scaled score of State B; State B bias score is calculated as the inverse. A positive State *A* bias score reflects a predicted higher likelihood to occupy that conformation. By default we refer to the bias score for a protein as the bias score towards the active conformation or the conformation of interest.

Mutations were filtered by selecting those with an individual scaled score above zero on at least one structure, as low scoring mutations on both structures were often low expression/low activity. The predicted State *A* biased mutants were assigned as the top 2.5% of filtered mutants with the highest State A bias score. The predicted State *B* mutants were assigned as the 2.5% of filtered mutants with the highest State B bias score. The neutral set of mutations (middle population in figures) was selected as the 95% confidence interval of unfiltered mutations based on assay score, representing the general distribution of mutant activities with outliers removed.

For all datasets analyzed in **Figures 2** and **5**, PDB files were downloaded from the ProteinDataBank (listed at end of Methods). Corresponding deep mutational scanning data was sourced from previous studies of those respective proteins. Any modified amino acids (e.g. pTyr, Sec) not included in the ProteinMPNN token alphabet were replaced by their base amino acid in the same orientation (e.g. Tyr, Cys).

DMS datasets were preprocessed to remove any missing values. In the case of growth assays where data was present for the detection of mutants before selection (MurA/FabZ), mutants that were detected but failed to grow (resulting in a missing value) were assigned the minimal activity score present in the dataset. DMS data may have a different frame of reference than the sequences associated with each structure, so a reference sequence was constructed from each DMS dataset and aligned to the structures for the respective protein. Only mutations at positions present in both structures were retained. Mutational data was then analyzed against calculated CB bias scores.

For K-Ras analysis, due to the large percentage of residues involved in binding with effectors, mutations from known binding interface residues(3) were removed from analysis. Mutations at positions directly flanked by binding interface residues were also filtered out, as shown in **Figure S1A**. For SARS-CoV2, due to the homo-trimeric structure of spike protein, scores were calculated on all three chains and averaged to calculate the "Up" and "Down" conformation scores. For β2AR, due to missing segments on each chain, AlphaFold2 predictions for (filtered to plDDT > 70) were used to increase coverage of the protein for CB analysis.

For FabZ, both structures are from *H. Pylori*. For MurA, the loop open structure is from *E. Cloacae*. Since the deep mutational scanning data for these proteins is from *E. coli,* AlphaFold2 was used in single sequence mode with 48 recycle iterations, templated against the respective structure in the desired conformation, and the highest confidence structure selected by pLDDT. Resulting AF2 structures were verified to have close alignment with regards to conformational state with the template.

The statistical significance of differences between populations was calculated using the Student's unpaired t-test. RMSD structural plots were generated using the colorbyrmsd function in PyMOL. All visualizations of structures were also performed in PyMOL.

**Conformational Biasing (CB) workflow for Synthetic Binders.** All structures were downloaded from the ProteinDataBank. Each pair of Domain *A*/Domain *B* sequences was scored across every experimentally solved binder structure (n=10). A synthetic distribution of scores was generated by sampling sequences (n=10000) from the original distribution of amino acid frequencies(1) used to generate the binders, and then scoring all sequences against each structure. This synthetic distribution of scores was then used to calculate the Z-score per structure for each of the original binders. A higher Z-score thereby represents a better fit for the binding mode found in a given experimental structure. Scores for matched structure/sequence pairs were then analyzed against scores for non-matched structure/sequence pairs.

**Conformational Biasing (CB) workflow for LplA.** Structures for LplA (PDBs: 3A7R(9), 1X2G(10)) were downloaded from the ProteinDataBank. Modified amino acids in the 1X2G structure (selenomethionine) were converted to the base residue. The two structures were confirmed to have the same sequence. All point mutants for the sequence were generated, and mutant sequences were scored on both structures using ProteinMPNN, as discussed above.

The mutant sequences were first filtered by requiring the score on at least one structure to be greater than that of wild-type, a more stringent cutoff than what was used for the DMS data. Open-bias score was calculated as the difference between the CB score on the open structure and CB score on the closed structure; closed-bias score is equal to negative open-bias score. Afterwards, three subpopulations of mutants were selected for testing. Predicted neutral mutants scored highly on both structures. Predicted open-biased mutants were selected based on high open-bias score (cutoff of 5.61 open-biasing score), and predicted closed-biased mutants were selected for high closed-bias score on 3A7R (cutoff of 6.50 closed-biasing score). Each subpopulation represents approximately 0.5% of all possible point mutants on the LplA sequence. Gain-of-lysine variants were largely excluded to avoid the possibility of novel mechanisms of auto-labelling activity, which might lead to inaccurate measurement of the promiscuous activity. 1 closed variant (V316P), and three neutral variants (N109A, N163A, and N163E), failed to clone. In total 31 open-biasing, 32 closed-biasing, and 36 neutral mutants were cloned for testing in mammalian cells.

**Benchmarking CB-predicted variants with BioEmu.** For SARS-CoV2 analysis spike protein analysis, seven up-biased and down-biased predicted variants were selected. These variants were filtered to only include those that had the expected corresponding change in ACE2 binding (Up = increased, down = decreased) as well as to be not located in the ACE2 binding interface. Due to the large size of the spike protein, only a subset of the structure was simulated (residues 297-712), which includes three well-folded domains including the RBD. For each variant, the variant sequence was simulated by running BioEmu with default parameters (n=4000), and then filtered to remove unphysical samples. One "Up" variant (F515L) and two "Down" variants (R454D, L425N) had <5% of samples pass filtering, and thus were removed from further analysis. The angle between the RBD domain and other two domains was calculated using PyMOL, and compared to the native angle in the "Up" and "Down" conformations. For each variant, a two-component gaussian mixture model was fit, and used to assign samples as in either an “up”-like or “down”-like conformation, which is shown in **Figure S11**.

For LplA, variant sequences were sampled in the same manner (n=500 for single mutants, n=100 for double mutants). The NTD of each sample was aligned to the NTD of 3A7R (open) and 1X2G (closed), and the CTD RMSD was calculated to each of those structures. A sample with 3A7R-RMSD less than the mean, and 1X2G-RMSD greater than the mean was classified as open-like, and vice versa for assigning closed-like structures. Samples with high RMSD for both structures were considered to be similar to neither structure. BioEmu predictions for LplA were then compared to experimental tryptophan fluorescence data.

**Benchmarking CB with other inverse folding models.** For scoring β2AR variants, ESM-IF1, Frame2Seq, and ThermoMPNN were installed from Github and used as instructed in their respective manuscripts(11-13). For simplest comparison, the experimental structures only were used, without any in-filling of missing residues. The score was calculated in the same way as with ProteinMPNN for all variants. For LplA analysis, the other inverse folding models were applied the same way, and then correlated with experimental promiscuous labeling data.

**AFCluster Analysis of LplA.** AFCluster was run on the LplA sequence to provide a comparison to the CB workflow. Specifically, the ColabDesign implementation of AFCluster was run using default parameters (min. sequences/cluster = 3, models = 1, recycles = 3) on the wild-type *E. coli* LplA sequence. Structural similarity of clusters to open- and closed- conformations was done by calculating pairwise TM-score(14) as done in the AFCluster paper(15). For each point mutation, the mean TM-score across all clusters it was present on was calculated, with mutants having a high mean TM-score towards the open conformation being enriched in open conformation predicted clusters and vice versa for closed conformation predicted clusters. These mean values were then scaled to a mean of 0 and standard deviation of 1. We calculated a value for enrichment in open vs. closed clusters, similar to the prediction of fold-switching mutations for KaiB in the original paper, by taking the difference between scaled mean open TM-score vs. scaled mean closed TM-score. This value was then compared to our experimental data on LplA.

**Cloning and mutagenesis of LplA.** All mammalian expression constructs were prepared in the pCDNA3.1 vector. The mCherry-LplA expression construct was cloned by Gibson assembly. All point mutations were cloned by amplification of the plasmid using mutagenic primers to introduce point mutations at the site of a Gibson assembly junction. Bacterial expression plasmids were cloned similarly by Gibson assembly of the *E. coli* codon-optimized LplA gene.

**LplA, LAP, and E2p protein expression in HEK 293T cells.** 24 well plates were coated with 200uL human fibronectin (HFN) for 30 minutes at 37°C. HFN was aspirated, and then HEK293T cells were plated at 10^5 cells per well. Following 12-24h of cell growth to 60-80% confluency, transfections were performed. For each well, 500ng of the mCherry-LplA plasmid and 4uL of PEI transfection reagent were combined in 50uL of DMEM without FBS or antibiotics, incubated for 20 minutes at room temperature, and then added to each well of a 24-well plate dish. Cells were grown for 24 - 48 hours after transfection to allow for LplA expression, and mCherry signal was confirmed by imaging. For co-expression of LplA variants with E2p acceptor protein, transfections were performed with a 1:50 ratio of mCherry-LplA to E2p-YFP DNA. For co-expression of LplA variants with LAP-mCherry, transfections were performed at a 1:1 ratio of mCherry-LplA to LAP2-mCherry DNA.

**Flow cytometry assay for promiscuous labeling by LplA variants.** Following successful transfection of cells, reagents necessary for labeling were prepared. A 200mM working stock of the labeling probe Endo-BCN Pentanoic Acid (BCN) (Broadpharm, BP-24361) was prepared by dilution of solid BCN in DMSO. A labeling solution of 200uM BCN in DMEM+FBS+penicillin/streptomycin (1000x dilution) was prepared from this. Next, a stock solution of 1mM methyltetrazine-BODIPY (Conjuprobe, CP-4018) was prepared in DMSO, then diluted 5000x in complete growth media. Both labeling and BODIPY solutions were pre-warmed to 37°C, along with additional complete media for washes. The growth media was removed from the transfected HEK293T cells, after which 0.5 mL of BCN solution was added to each well. The cells were moved to a 37°C cell culture incubator for 5 minutes, and then labeling media was removed by aspiration and immediately replaced with warmed complete media. Three additional 5-minute washes with warmed complete media were performed to remove excess BCN substrate. Afterwards, the media was aspirated and replaced with 0.5mL/well of the fluorophore-click solution (200nM mTz-BODIPY) and cells were incubated at 37°C for 45 minutes. Media was then removed by aspiration and cells were quickly washed twice in pre-warmed complete media. Three additional 15-minute washes (15 min incubation between media addition and aspiration) in complete media were performed to remove unbound fluorophore. Once media from the final wash was removed, cells were incubated for 1 minute in 100uL of enzyme-free cell dissociation buffer at 37°C, and then lifted with an additional 100uL flow buffer (PBS + 3% FBS) and transferred to a 96-well plate. Cells were pelleted once by spinning at 400rcf for 5 minutes at 4°C, and then resuspended in flow buffer before analyzing on a BioRad ZE5 flow cytometer. BFP was detected using a 405 nm laser, with a 460/22 emission filter. mTz-BODIPY was detected using a 488 nm laser, with a 509/24nm emission filter. mCherry was detected using a 561 nm laser, with 615/24 nm emission filter. Forward and side scatter were detected using the 488 laser with a 488/10 emission filter. Each sample was agitated for 5s, and run with a flow rate of 1ul/s.

**Quantitation of flow cytometry data.** Single cells were gated-for by forward scatter/side scatter profiles as above. After gating, cells were plotted by mCherry against FITC channels, and a diagonal gate was used to capture the linear region for the mCherry:BODIPY curve (**Figure S6A**). Cells within this gate were analyzed for median BODIPY / mCherry signal to quantify the promiscuous labelling activity of each variant. All data was analyzed using the software package FlowJo.

**SDS-PAGE analysis of LplA sequence-specific and promiscuous activity**. HEK293T cells in a 24-well plate were transfected as described above. 24-48 hours after transfection, after validation of expression of transfected constructs, the cell growth media was removed and labeling was carried out by addition of pre-warmed labeling media (200uM BCN in complete DMEM) for 5 minutes at 37C. Labelling was quenched with 3x washes with ice-cold PBS.

To lyse cells for protein enrichment, RIPA buffer was supplemented with 1:100 dilutions of protease inhibitor cocktail (PIC) and phenylmethylsulfonyl fluoride (PMSF). Labeled cells were resuspended by pipetting in 100 µL of RIPA buffer, and cell suspensions were incubated on ice for 10 minutes at 4°C. Following lysis, cell debris was removed by centrifuging the lysate at 17,900 × g for 10 minutes at 4°C. The supernatant was collected for subsequent analyses.

BCN-labeled proteins were further labelled via the inverse-electron demand Diels-Alder Cycloaddition (IEDDAC) reaction of BCN with methyl-tetrazine (mTz). For in-gel fluorescence analysis, either mTz-BODIPY, or mTz-JF647, were added to the clarified lysate at a final concentration of 4uM and allowed to react for 2 hours at 4°C with vortexing at 850rpm. Unreacted methyl-tetrazine dye was then removed from the lysate by addition of 1ul of TCO-agarose per 50ul of lysate, and incubation for 1 hour at 4°C with vortexing at 850rpm.

Clicked and purified lysates were mixed 5:1 with protein loading buffer and mixed thoroughly. Importantly, samples were not boiled, to retain fluorescence of the fluorescent protein tags mCherry and YFP. 6ul of each lysate was loaded per well of gradient SDS-PAGE gels (4-12%) (BioRad), and run in SDS running buffer (Tris-Glycine) for 45 minutes at 180V. Gels were then washed briefly in milliQ water, and then imaged on a Typhoon biomolecular imager. mTz-BODIPY was imaged using a 480nm laser, with a 520nm emission filter. YFP and mCherry were imaged using a 532nm laser, with 610nm emission filter. JF-647 was imaged with a 633nm laser, using a 670nm emission filter.

IGF data was analyzed and quantified using FIJI. Labelling and expression were background-subtracted, and the labelling was normalized to expression level for each variant to calculate the expression-normalized activity. Then the expression-normalized activity of each variant is divided by the expression-normalized activity of the template W37V to calculate the relative activity for each variant.

**Confocal microscopy.** 48 well glass-bottom plates were coated with 100uL human fibronectin (HFN) for 30 minutes at 37°C. HFN was aspirated, and then HEK293T cells were plated at 4.5E4 cells per well. Following 12-24h of cell growth to 60-80% confluency, transfections were performed. For each well, 250ng of the mCherry-LplA plasmid and 2uL of PEI transfection reagent were combined in 25uL of DMEM without FBS or antibiotics, incubated for 20 minutes at room temperature, and then added to each well of the 48-well plate dish. Cells were grown for 24 - 48 hours after transfection to allow for LplA expression.

Labelling was carried out as in the flow-cytometry assay. The cell growth media was removed and labeling was carried out by addition of pre-warmed labeling media (200uM BCN in complete DMEM) for 5 minutes at 37C, and then immediately replaced with warmed complete media. Three additional 5-minute washes with warmed complete media were performed to remove excess BCN substrate. Afterwards, the media was aspirated and replaced with 0.5mL/well of the fluorophore-click solution (200nM mTz-BODIPY) and cells were incubated at 37°C for 45 minutes. Media was then removed by aspiration and cells were quickly washed twice in pre-warmed complete media. Three additional 15-minute washes (15 min incubation between media addition and aspiration) in complete media were performed to remove unbound fluorophore. Once media from the final wash was removed, cells were washed 3x in PBS, and treated with 4% paraformaldehyde for 15 minutes at room temperature. Cells were again washed 3x in PBS, and imaged using an Olympus APX100 microscope with a 20x objective. mTz-BODIPY was detected using a 480 nm laser, with a 520nm emission filter, and mCherry was detected using a 532 nm laser, with 610 nm emission filter. Images were analyzed using FIJI.

The following combinations of laser excitation and emission filters were used for various fluorophores: Bodipy (491 laser excitation; 528/38 emission), mCherry (561 laser excitation; 617/73 emission). Acquisition times ranged from 100 to 500 ms. All images were collected using SlideBook (Intelligent Imaging Innovations) and processed using FIJI/ImageJ.

**Expression and purification of LplA from *E. coli*.** Chemically competent BL21 (DE3) pLysS *E. coli* cells were thawed on ice and then transformed with plasmids encoding LplA variants following the producer's recommended transformation protocol. Transformed cells were plated onto LB-ampicillin agar plates and incubated overnight at 37 °C. After incubation and observation of colonies, 5–10 colonies were picked and inoculated into 5 mL of LB-ampicillin medium and then shaken at 220RPM at 37 °C for over 6 hours until the culture became visibly cloudy. The 5 mL overnight culture was added to 500 mL of LB-ampicillin in a 2-liter flask. The culture was shaken at 37 °C until an A600 of 0.5 was reached, approximately 2–3 hours depending on the starting culture. The flask was then moved to a room-temperature shaker for cooling, and isopropyl-β-D-thiogalactopyranoside (IPTG) was added to a final concentration of 100 µg/mL. Induction was performed by shaking the flask at 220RPM overnight at room temperature (~8 h).

Cells were pelleted by centrifugation at 5,000 × g for 10 min at 4 °C. The supernatant was discarded, and the bacterial pellet was kept on ice. For each sample, 100 µL of 100 mM PMSF and 100 µL protease inhibitor cocktail were added to 20 mL Bacterial Protein Extraction Reagent (B-PER). The pellet was lysed by thoroughly resuspending in 10 mL of the prepared B-PER. After thorough resuspension, an additional 10 mL of the prepared B-PER was then added to ensure thorough suspension. The suspension was gently agitated for 10 min at 4 °C. The lysate was transferred to high-speed centrifuge tubes and centrifuged at maximum speed for 60 min at 4 °C. The resulting supernatant, containing the cell lysate, was collected in a 50 mL conical tube.

Nickel-NTA resin for purification was prepared by loading approximately 1 mL of packed resin into a Poly-Prep column, followed by washing with five column volumes of ice-cold nickel-binding buffer. 20 mL of ice-cold nickel-binding buffer was added to the cell lysate, and the prepared resin was transferred to the lysate tube. The mixture was gently agitated for 20 min at 4 °C to facilitate His-tagged protein binding to the resin. The cell lysate/resin mixture was loaded onto a Poly-Prep column, avoiding the column drying out. The resin was washed with ten column volumes of nickel-NTA wash buffer to remove non-specifically bound proteins. Bound protein was eluted using 3 mL of nickel-NTA elution buffer. Elution fractions were collected in 500 µL volumes and checked for protein content by measuring the A280 absorbance.

As an optional additional step, for tag-less preparation, positive fractions as defined by A280 absorbance were pooled, and DTT and glycerol were added to a working concentration of 2 mM and 5% v/v respectively. His-TEV protease (Berkeley QB3 macro-lab) (1:50 molar ratio) was added to the eluate for cleavage, and the sample was incubated overnight at 4 °C. The eluate was then diluted 20-fold to achieve DTT and imidazole concentrations of <0.2 mM and <5 mM, respectively. Nickel-NTA resin (~1 mL packed per ~5 mg protein) was prepared and washed, then used to bind the reaction mixture for 20 min with gentle agitation at 4 °C. The suspension was loaded onto the column, and the flow-through, containing tag-less LplA, was collected. The resin was washed with one column volume of binding buffer, and the flow-through was again collected and combined.

DTT and glycerol concentrations were re-adjusted to 2 mM and 5% v/v, respectively. The protein solution was concentrated to <500 µL using a protein concentrator (Amicron Ultra, MW Cutoff 10000Da), maintaining a concentration below 10 mg/mL of protein to minimize aggregation. The sample was centrifuged at 10,000 × g for 10 min, and the supernatant was loaded onto a pre-equilibrated SEC column. A280 positive fractions were collected, and again the protein solution was concentrated using a protein concentrator (Amicron Ultra, MW Cutoff 10000Da), maintaining a concentration below 10 mg/mL of protein to minimize aggregation. Protein concentration was then quantified using a DTT-compatible BCA assay, and protein samples were aliquoted, flash-frozen in liquid nitrogen, and stored at –80C until use.

**Synthesis and characterization of lipoyl-AMS.** A glass vial equipped with a magnetic stir bar was charged with 18.86mg of 2,5-dioxopyrrolidin-1-yl (R)-5-(1,2-dithiolan-3-yl)pentanoate (62.17 µmol), 20 mg 2',3'-isopropylidine sulfamoyladenosine (ChemPartner, 51.81 µmol), and 22 µL 1,8-diazebicycloundec-7-ene (142 µmol) dissolved in 2 mL N,N-dimethylformamide. The mixture was stirred at room temperature for 16 hr. After 16 hours, the mixture was diluted with 10 mL H2O then extracted three times with 30 mL ethyl acetate. The combined organic washes were dried with sodium sulfate and the volatiles removed via rotary evaporation. The resulting solution solid was stirred in 2 mL 80% TFA in water for 90 min. The TFA was then removed by rotary evaporation and the remaining solution was loaded onto an HPLC (Waters). Purification of the desired product was carried out by successive 5-mL elutions of 50-100% acetonitrile in H2O, 0.05% TFA (v/v). The aqueous solution was freeze-dried to give 6.71 mg of white solid (over yield 24.2 %).

$^1$**H NMR** (500 MHz, DMSO-d6) δ 8.35 (s, 1H), 8.22 (s, 1H), 5.88 (d, J = 7.0 Hz, 1H), 4.50 – 4.34 (m, 3H), 4.16 – 4.08 (m, 2H), 3.12 – 3.01 (m, 3H), 2.31 (m, 1H), 2.18 (m, 2H), 1.77 (m, 1H), 1.57 – 1.36 (m, 4H), 1.28-1.15(m, 2H); **LC-MS** m/z: calculated for [M+H]+ 535.11, Found 535.14

**Tryptophan fluorescence assay.** Purified LplA variants from Nickel-NTA purification above were thawed on ice. For the tryptophan fluorescence assay, each protein was diluted to a final concentration of 10 µM in the assay buffer (see below). Protein concentration was quantified by BCA assay (ThermoFisher, 23235), followed by SDS-PAGE and Coomassie to normalize the protein concentration between samples. SEC buffer was supplemented to the dilution series such that the final buffer composition for all variants was equivalent.

To establish baseline fluorescence, 30 µL of each protein solution at 10 µM was transferred into individual wells of a 384-well white microplate (Costar). Experiments were performed with three technical replicates each. Baseline fluorescence measurements were recorded using a Tecan plate reader (Molecular Devices) at room temperature, with excitation and emission wavelengths of 290 nm and 340 nm, respectively. Z-position and Gain were calculated based on the first well containing protein sample. Measurements were recorded every ~30 seconds for over 20 minutes to establish a baseline measurement of intrinsic fluorescence.

Following the baseline readings, 3 µL of Lipoyl-AMS in assay buffer was added to each well, containing 1mM Lipoyl-AMS, or 500uM Lipoyl-AMS (each prepared from a 10 mM stock). This addition brought the total well-volume to 33 µL, with a final L-AMP concentration of 100 µM, or 50uM. Immediately after the addition of the assay stock, the plate was read again using the same excitation and emission settings to monitor the fluorescence drop. Measurements were taken every ~30 seconds for 30 minutes after L-AMP addition, such that equilibrium was visibly reached for all protein samples.

**Tryptophan fluorescence data analysis.** Pre and post lipoyl-AMS addition fluorescence intensity was measured by averaging across measurements taken over the course of 10 minutes prior to Lipoyl-AMS addition, and averaging across measurements taken over the course of 15 minutes after fluorescence had stabilized following Lipoyl-AMS addition. The change in fluorescence for each sample was calculated by dividing these values (fluorescence after lipoyl-AMS addition / baseline fluorescence before lipoyl-AMS addition). Each variant was measured in triplicate, and could then be compared to the template, W37V, using a student's t-test (two-tailed, two-sample equal variance, $p<0.01$). Next, the triplicate measurements of change in fluorescence were averaged, and the standard deviation calculated for each variant. Then, each was divided by the change in fluorescence of the template W37V, to determine the change in tryptophan fluorescence upon Lipoyl-AMS addition, relative to W37V. Error bars were calculated by propagating the standard deviation of the variant in question, and the standard deviation of the template W37V, following the standard rules of propagation of error for division of values.

**Site-specific protein labeling in cells with LplA.** HEK 293T cells (for NLS-BFP) and HeLa cells (for HA-beta-actin) were plated at 70% confluency in glass-bottomed 24-well plates. 8 hours after plating, cells were transfected using 1 µL Lipofectamine 3000 and 1 µL P3000 with LAP2-tagged constructs and either W37V or T57I mCherry-LplA. 334 ng of LAP2 plasmid was used and 166 ng of LplA plasmid was used. Twenty hours post-transfection, LplA labeling was performed using 200 µM BCN in DMEM + 10% FBS for 10 minutes at 37°C for NLS-BFP, and 20 µM coumarin-AM2 in serum-free DMEM supplemented with 0.1% Pluronic F-127 for 10 minutes at 37°C for HA-beta-actin. All cells were then washed three times with warm media. For NLS-BFP, an in-cell Click labeling was performed using 500nM methyltetrazine-BODIPY in HBSS, followed by three washes with warm HBSS. NLS-BFP cells underwent an additional 1-hour washout in DMEM + 10% FBS before live cell imaging. For HA-beta-actin, cells were fixed with 4% paraformaldehyde for 15 minutes at room temperature, washed three times with DPBS, permeabilized with 0.25% Triton X-100 for 10 minutes, washed three times with DPBS, then blocked overnight at 4 °C in 1% BSA in DPBS. Cells were then stained with Rabbit anti-HA antibody (1:500) for 1 hour at room temperature, washed three times, then stained with Goat anti-Rabbit AlexaFluor 647 antibody (1:1000) for 1 hour, and washed again before imaging.

All cells were imaged by confocal microscopy using a Zeiss AxioObserver inverted microscope equipped with a Yokogawa spinning disk confocal using a 100x oil-immersion objective. Laser excitation and emission filters for fluorophores are as follows: BFP/Coumarin (381 laser, 445 emission), BODIPY (491 laser excitation; 528/38 emission), mCherry (561 laser excitation; 617/73 emission), Alexa Fluor 647 (647 excitation; 680/30 emission). Acquisition times ranged from 100 (mCherry, AF647) to 500ms.

**Modeling of LplA reaction progress.** LplA reactions were modeled in Python assuming Michaelis-Menten kinetics and with the following initial conditions:

Enzyme concentration: approximate as 10 uM
Cognate substrate concentration: approximate as 40 uM
Non-cognate substrate concentration: approximate as 1 mM
Cognate substrate $K_M$: 1 uM
Non-cognate substrate $K_M$: approximate as 8 mM
Cognate substrate $K_{Cat}$: 0.1 $s^{-1}$
Non-cognate substrate $K_{Cat}$: approximate as 0.02 $s^{-1}$ to 0.001 $s^{-1}$

It was assumed that shifting the $K_{Conf}$ for open and closed-biased LplA variants would result in a 4x decrease or increase in $K_M$ for open and closed biased variants, respectively. To change the model such that both substrates were above their respective $K_M$ values, the non-cognate substrate $K_M$ was lowered to 80 uM. To change the model such that both substrates were below their respective $K_M$ values, the cognate substrate concentration was lowered to 400 nM. Non-cognate substrate $K_{cat}$ is varied from model to model to scale non-cognate rates to better visualize non-cognate substrate reaction curves.

**SEC-SAXS experiments and analyses.** The SEC-SAXS experiments were conducted at the Stanford Synchrotron Radiation Light Source (SSRL) Bio-SAXS beamline 4-2(19). SEC-SAXS data were collected at High-Resolution mode using a Superdex 75 Increase 3.2/300 column (Cytiva) using DTT-supplemented SEC running buffer (50 mM Tris pH 8.0, 150 mM NaCl, 5mM DTT). Protein samples were prepared in this buffer at 4.3mg/ml for the wild-type LplA and 7.1 mg/ml for the A48N mutant. Proteins were purified without N-terminal His-tag, following the tag-less purification method as described above. SAXS samples were prepared by addition of 1mM lipoic Acid or 1mM lipoyl-AMP to these proteins and incubation for 20 minutes at 4C prior to SEC-SAXS. 500 images were acquired with 2-second exposure every 5 seconds at a flow rate of 0.05 ml/min. To minimize the sample cell fouling problem, the x-ray shutter was closed after the 100th image (blank data collection) and the sample cell was immediately washed in preparation for the following fractions of interest. The shutter was then re-opened to resume the image acquisitions at the fractions. Evolving factor analysis was applied to the WT+lipoyl-AMP data set to improve the signal-to-noise ratio using the program *BioXTAS RAW(20)*. Data reduction and initial analyses were performed using the BL4-2 automated SEC-SAXS data processing and analysis pipeline, *SECPipe* (https://www-ssrl.slac.stanford.edu/smb-saxs/node/1860). It implements the programs *SASTOOL* (https://www-ssrl.slac.stanford.edu/smb-saxs/node/1914) and *ATSAS AUTORG(16)*. The data were plotted as I(q) versus q, where q = 4πsin(θ)/λ, 2θ is the scattering angle, and λ is the wavelength of the X-ray. After careful manual inspection, a total of 5 images were selected to generate the averaged profile for further analysis.

The program *Oligomer* was used to estimate the volume fractions of open and closed conformations(17). Atomic models of the open and closed conformations were generated from the existing structures: PDB 1X2H Chain A and 3A7R comprise the closed and open monomers, respectively. After observing a likely dimeric fraction for the open-conformation of A48N + lipoyl-AMP, we used AlphaFold3 to predict the structure of the dimeric A48N LplA in complex with 2 AMP ligands, and the resulting structure was used as the atomic model for the open-dimeric conformation for estimation of volume fractions. The program *GNOM* was used for the indirect Fourier transform to estimate the distance distribution function *P*(r) (18). *Ab initio* electron density maps of the wild-type and A48N mutant were calculated using the program *DENSS(19)*, with a final map calculated by averaging 20 independent runs. Experimental and analytical details are summarized below.

| | WT+LA | WT+LAMP | A48N+LA | A48N+LAMP |
|---|---|---|---|---|
| ***Data collection parameters*** | | | | |
| Instrument | SSRL BL4-2 | | | |
| Type of Experiment | SEC-SAXS | | | |
| Beam Current (mA) | 500 | | | |
| Defining slits size (H mm × V mm) | 0.3 × 0.30 | | | |
| Detector distance (m) | 1.1 | | | |
| Detector | Pilatus3 X 1M | | | |
| Beam energy (keV) | 11.0 | | | |
| *q* range ($Å^{-1}$) | 0.009–0.73 | | | |
| Sample cell | Quartz capillary (*ID*=~1.3mm) | | | |
| Temperature (K) | 295 | | | |
| Exposure time/frame (s) | 1 | | | |
| Frames per SEC-SAXS data set | 500 | | | |
| Number of blank images used for averaging | 50 | | | |
| Number of sample images used for averaging | 5 | | | |
| Image numbers used for averaging | N/A | 300 - 304 | 315 - 319 | 280 - 284 |
| SEC column | Superdex 75 Increase 3.2/300 | | | |
| HPLC flow rate (mL/min) | 0.05 | | | |
| Sample concentration (mg/ml) | 4.3 | 4.3 | 7.1 | 7.1 |
| SEC injection volume (μL) | 100 | 30 | 100 | 50 |
| Buffer | SEC Buffer with 5mM DTT | | | |

| | | | | |
|---|---|---|---|---|
| ***Software employed*** | | | | |
| Primary data reduction | *SasTool/SECPipe* | | | |
| Data processing | *PRIMUS* | | | |
| P(r) analysis | *GNOM* | | | |
| Volume fraction estimation | *Oligomer* | | | |
| *ab initio* modeling | *DENSS* | | | |
| ***Structural parameters*** | | | | |
| *Guinier analysis* | | | | |
| *I*(0) (*arbitrary unit*) | 2205.03 ± 4.03 | 334.58 ± 0.38 | 241.81 ± 0.34 | 555.30 ± 0.67 |
| $R_g$ (Å) | 22.79 ± 0.06 | 22.87 ± 0.04 | 22.98 ± 0.05 | 32.62 ± 0.06 |
| $qR_g$ range | 0.35 – 1.29 | 0.32 – 1.30 | 0.29 – 1.30 | 0.65 – 1.30 |
| *P(r) analysis* | | | | |
| *I*(0), Guinier (*arbitrary unit*) | 2217 | 334.5 | 239.7 | 560.3 |
| $R_g$ (Å), Guinier | 23.04 | 22.84 | 22.86 | 33.79 |
| *I*(0), *P(r)* (*arbitrary unit*) | 2217 | 334.5 | 239.7 | 560.3 |
| $R_g$ (Å), *P(r)* | 23.03 | 22.84 | 22.86 | 33.85 |
| $D_{max}$ (Å) | 74.61 | 74.32 | 73.00 | 115.90 |
| q range (Å$^{-1}$) | 0.0156 – 0.351 | 0.0140 – 0.350 | 0.0125 – 0.348 | 0.011 - 0.245 |
| Porod volume estimate (Å$^3$) | 70187 | 67965 | 60915 | 85419 |
| ***Oligomer*** | | | | |
| $q_{max}$ (Å$^{-1}$) | 0.4 | 0.4 | 0.4 | 0.4 |
| $c^2$ | | | | |
| ***DENSS*** | | | | |
| Mode | Slow | | | |
| Symmetry | P1 | | | |
| Number of reconstructions | 20 | | | |
| $q_{max}$ (Å$^{-1}$) | 0.7268066 | 0.7268066 | 0.7268066 | 0.7268066 |
| $c^2$ | | | | |
| Resolution (FSC = 0.5) (Å) | 35.783 | 36.530 | 33.425 | 34.924 |

| Sample Conditions | | | Protein parameters | | | Protein Concentration | |
|---|---|---|---|---|---|---|---|
| Sample # | Protein | Condition | MW | Ext. Co | Abs. | mg/ml | uM |
| 1 | A48N | LA-AMS, 1% DMSO | 38025.97 | 47440 | 1.248 | 5.5 | 145 |
| 2 | WT | LA-AMS, 1% DMSO | 37982.94 | 47440 | 1.249 | 4.75 | 125 |
| 3 | C55M$^{V}$ | LA-AMS, 1% DMSO | 37923.91 | 41940 | 1.106 | 5.09 | 134 |
| 4 | W37V | LA-AMS, 1% DMSO | 37895.86 | 41940 | 1.107 | 1.13 | 30 |
| 5 | T57I$^{V}$ | LA-AMS, 1% DMSO | 37907.91 | 41940 | 1.106 | 1.7 | 45 |
| 6 | A48N$^{V}$ | LA-AMS, 1% DMSO | 37938.88 | 41940 | 1.105 | 2.26 | 60 |
| 7 | E265R$^{V}$ | LA-AMS, 1% DMSO | 37922.93 | 41940 | 1.106 | 1.98 | 52 |
| 8 | A48N | Buffer | 38025.97 | 47440 | 1.248 | 5.5 | 145 |
| 9 | WT | Buffer | 37982.94 | 47440 | 1.249 | 4.75 | 125 |
| 10 | C55M$^{V}$ | Buffer | 37923.91 | 41940 | 1.106 | 5.09 | 134 |
| 11 | W37V | Buffer | 37895.86 | 41940 | 1.107 | 1.13 | 30 |
| 12 | T57I$^{V}$ | Buffer | 37907.91 | 41940 | 1.106 | 1.7 | 45 |
| 13 | A48N$^{V}$ | Buffer | 37938.88 | 41940 | 1.105 | 2.26 | 60 |
| 14 | E265R$^{V}$ | Buffer | 37922.93 | 41940 | 1.106 | 1.98 | 52 |

**Reagents used in this study.** Endo-BCN Pentanoic Acid (BCN) (Broadpharm, BP-24361), Methyltetrazine-BODIPY (Conjuprobe, CP-4018-1mg), JF647-methyltetrazine (gift from Pratik Kumar and Luke Lavis, Tocris, 7279), Lipoyl-AMP (synthesized by Daniel Liu).

**Buffers:**

- *Nickel-NTA Binding Buffer*: Prepared by mixing 50 mM Tris base and 300 mM NaCl, adjusted to pH 7.8 with 1 M HCl and stored at 4 °C.
- *Nickel-NTA Wash and Elution Buffers*: Prepared by adding 1 M imidazole to binding buffer. For wash buffer, 10 mM imidazole was used, and for elution buffer, 100 mM imidazole.
- *SEC Buffer*: Prepared with 40 mM Tris (pH 7.5), 1 mM DTT, 0.5 mM EDTA, and 150 mM NaCl.
- *Tryptophan Fluorescence Assay Buffer:* Dulbecco's Phosphate-Buffered Saline (DPBS), 1 mM Dithiothreitol (DTT), 5 mM Magnesium Chloride ($MgCl_2$), and 0.05% (v/v) Tween-20 detergent, 5% (v/v) glycerol.

**Genetic constructs used in this study**

| Construct | Vector | Expression | Description | Used in |
|---|---|---|---|---|

| | | | | |
|---|---|---|---|---|
| His6-mCherry-Flag-LplA | pcDNA3 | Mammalian | Constructs used for LplA-catalyzed labeling in mammalian cells. LplA is expressed as a fusion with mCherry. | Figure 4 |
| LAP2–BFP-CAAX | pcDNA3 | Mammalian | Plasmid used to express LplA's site-specific substrate LAP2 (13-amino acid acceptor peptide) in mammalian cells. CAAX targets it to the inner leaflet of the plasma membrane. | Figure 4 |
| LAP2–BFP-NLS | pcDNA3 | Mammalian | LAP2 fused to BFP and NLS, a nuclear localization signal. Imaging shows primarily nucleolar localization. | Figure 4 |
| HA-LAP2-actin | peGFP | Mammalian | LAP2 fused to HA epitope tag and beta-actin | Figure 4 |
| His6-E2P-YFP | pcDNA3 | Mammalian | Plasmid used to express E2p, a site-specific acceptor protein for LplA, in HEK293T cells. Used to evaluate specific labeling activity of LplA variants. | Figure S6F |
| His-TEV-LplA | pET | Bacterial | Plasmid used for bacterial expression of LplA variants. TEV cleavage site enables removal of N-terminal His6 tag. | Figure 3 |

**PDB structures used in this study**

- LplA – Open Conformation (Lipoyl-AMP bound): 3A7R; Closed Conformation (Apo): 1X2G, Closed Conformation (Lipoic Acid bound): 1X2H; Open Conformation in complex with target protein ApoH and Octyl-AMP: 3A7A.
- K-Ras – State 1: 8T71, State 2: 6XHB
- SARS-CoV2 – Down: 7XIX, Up: 7XO8
- β2AR – Inactive: 3SN6, Active: 2RH1
- Synthetic Binders: 8DA3, 8DA4, 8DA5, 8DA6, 8DA7, 8DA8, 8DA9, 8DAA, 8DAB, 8DAC
- Src Kinase – Inactive: 2SRC, Active: 1Y57
- B-Raf – Monomeric: 7MFD, Dimeric: 7MFF
- FabZ – Closed: 2GLL, Open: 4ZJB
- MurA – Loop Open: 1NAW, Loop Closed: 1UAE
